# Fish as model systems to study epigenetic drivers in human self-domestication and neurodevelopmental cognitive disorders

**DOI:** 10.1101/2022.02.24.481892

**Authors:** Dafni Anastasiadi, Francesc Piferrer, Maren Wellenreuther, Antonio Benítez Burraco

## Abstract

Modern humans exhibit phenotypic traits that are shared across independent domestication events, suggesting the human self-domestication hypothesis. Epigenetic changes may facilitate early self-domestication in humans, since they can be the first layer of response to a novel environment. Here, we argue that fish provide model systems to study epigenetic drivers in human self-domestication. To do this, we compare genes that carry epigenetic changes in early domesticates of European sea bass with 1) anatomically modern humans and 2) neurodevelopmental cognitive disorders with abnormal self-domestication traits, i.e., schizophrenia, Williams syndrome and autism spectrum disorders. We found that genes with epigenetic changes in fish and in modern vs ancient humans were shared and were involved in processes like limb morphogenesis and phenotypes like abnormal snout morphology and hypopigmentation. Moreover, early domestication in fish and neurodevelopmental cognitive impairment in humans affected paralogue genes involved in processes such as neural crest differentiation and ectoderm differentiation. We conclude that parallel epigenetic changes may occur at the initial steps of domestication in absence of deliberate selection in phylogenetically distant vertebrates. These findings pave the way for future studies using fish as models to investigate epigenetic changes as drivers of human-self domestication and as triggers of cognitive disorders.

## 1. Introduction

Domestication is a multifactorial process that is induced and maintained by human activity and human-generated environments. Historically and contemporary, this process has affected the evolutionary trajectories of several economically and culturally important vertebrate species. New phenotypic traits emerge repeatedly in independent vertebrate domestication events, even at the early stages of living in a human-made environment prior to deliberate selection; a phenomenon characterized as the *domestication syndrome* [1]. The domestication syndrome has been predominantly described in mammals, likely due to the large number of mammalian domesticates with a long domestication history, sometimes dating back millennia (e.g. dogs). Phenotypic traits of the domestication syndrome include a decreased size of the brain, heart and teeth, vertebrae variability, caudal vertebrae changes, shorter muzzle, more frequent estrous cycles, floppy ears, curly tail and hair, and depigmentation [2,3]. These traits have all been considered to have arisen as by-products of selection for increased tameness. Since these traits are associated with the final sites of migration of neural crest cells, mild developmental deficits affecting their development, migration or differentiation have been suggested as underlying mechanisms of the domestication syndrome, termed the *neural crest cell hypothesis* (NCCH) [1,4].

Modern humans, compared to extant apes and extinct hominins, exhibit phenotypic traits similar to those of other domesticated vertebrates, suggesting these may have also been produced as a by-product of selection for reduced aggression and increased sociality [5–7]. This is called the human self-domestication hypothesis [5,7]. Domestication syndrome-like morphological traits in anatomically modern humans (AMH) include a decreased brain and teeth size, facial robusticity, and sexual dimorphism, as well as neoteny [5,6,8]. Behavioral traits include reduced aggression, increased sociability, prolonged playing behavior, and overall more flexible social skills [5,6,8]. For understanding and evaluating the human self-domestication hypothesis, we need to distinguish between deliberate selection for improved traits, as occurred in, e.g., agricultural animals, and non-deliberate selection for prosociality arising from adaptation to novel environments, as expected for species hypothesized to have gone through a self-domestication process. The latter should be seen through the lens of domestication being a multi-stage process, where non-deliberate selection arises in response to the new selective environment, e.g. often involving a lack of predators and an increase in food availability [6], changed environmental conditions [9] or the colonization of new environments [10], which are all known factors promoting prosocial behavior. Empirical support for the human self-domestication hypothesis is challenging to obtain, nevertheless, comparative genomics have provided tentative support for it [11]. Recent results of an elegant study by Zanella et al. [12] using a molecular genetics approach are consistent with both the NCCH and the process of human self-domestication, specifically with regards to changes in the skull and the face [12,13].

Neurodevelopmental disorders in humans characterized by social and cognitive impairments may abnormally present traits of the domestication syndrome and thus may be linked to altered self-domestication. This is consistent with the view of self-domestication as a variable phenotype in human species, e.g. [14], with this variability depending on genetic and environmental factors. People with autism spectrum disorders (ASD) and schizophrenia (SZ) exhibit abnormal aggressive behaviour, abnormal responses to social cues, as well as tooth, ear and facial anomalies [15,16]. In ASD, increased head and brain size, and generalized overgrowth are also present, while in SZ, decreased brain volume and reproductive dysfunctions occur [15,16]. Accordingly, these two cognitive disorders can be regarded as “less self-domesticated” and “more self-domesticated” phenotypes, respectively [15,16]. Also Williams-Beuren syndrome (WS), caused by the hemideletion of 28 genes, is a clear example of a “more self-domesticated” phenotype [8,12]. People with WS show hypersociability, decreased aggression, reduced head and brain size, pointy ears, small teeth and jaws, depigmentation and accelerated sexual maturity [8]. Zanella et al. [12] used cell lines derived from WS subjects to establish the molecular links of morphological and behavioural domesticated traits in humans with neural crest development and migration. Therefore, cognitive disorders and the gene networks associated with them may be used as models for further testing of the human self-domestication hypothesis.

Domestication is a process of adaptation to a new selective environment, and has been considered as likely involving epigenetic changes [17–21]. Epigenetic mechanisms offer a way for novel phenotypes to emerge rapidly in response to environmental changes and to prime the offspring, when inherited, to face environments based on the parental experience [22–24]. In the first stages of domestication, which coincides with the emergence of domestication syndrome traits, epigenetic changes established during early development can regulate gene expression in the neural crest, and be maintained throughout adulthood and inherited to the offspring. Multigenerational epigenetic inheritance is ubiquitous in diverse animal species (see [25] for review). Persistence of the domestication environment, together with the stability and small effect of epigenetic changes in mild developmental deficits of the neural crest, are expected to accelerate adaptation [26]. After several generations, epigenetic changes could be genetically assimilated as genetic variants [21,27,28], hardwiring these changes. Partial evidence for this process comes from studies on mammals (dogs-wolves [19]), birds (red jungle fowl-modern chickens [29]), and fish species. The same process could be hypothesized to account for the first steps of human self-domestication, as most differences between extinct hominins and AMHs are epigenetic by nature, having impacted on features that are related to the domestication syndrome, particularly those impacting the face [30].

Domestication of fish species has a distinct history from terrestrial vertebrates [31], although it is scientifically considered to represent a similar process [32]. Until the 20^th^ century the majority of seafood has relied on wild animal captures, with few exceptions like the common carp (*Cyprinus cyprio*) in China ∼8000 years ago or Nile tilapia (*Oreochromis nilocitus*) in Egypt ∼3500 years ago [32,33]. In the last century, domestication of aquatic species has expanded rapidly, with an estimated number of 368 vertebrates that have been domesticated for aquaculture, teleost fish, frogs and reptiles [34]. Nevertheless, the majority of species are at the early stages of domestication, without closed life cycles in captivity and in the absence of deliberate selection for specific traits [34]. Nonetheless, in parallel with the domestication process, phenotypic traits involving the domestication syndrome, with changes in growth, reproduction, morphology, pigmentation and behaviour, have become manifested in domesticated fish [35–37]. Furthermore, sequencing of fish genomes has revolutionized vertebrate comparative genomics and has greatly contributed to our understanding of selection targets, evolutionary changes and speciation. Subsequently, fish have been suggested to serve as suitable models for human biomedical research [38,39]. Recently, epigenetic patterns emerging during the first stages of domestication, in the absence of genetic differences, have been studied in salmonids [40,41], European sea bass (*Dicentrarchus labrax*) [35], Nile tilapia (*Oreochromis niloticus*) [42,43] and grass carp (*Ctenopharyngodon idellus*) [44]. These epigenetic patterns of domestication are present in the sperm of several species, i.e. salmonids [41,45–47], showing the potential of intergenerational transfer, while in the European sea bass ∼20% are found in early embryos, showcasing the importance of developmental aspects during early domestication [35]. Taken together, 1) the recent domestication events in fish, 2) the high degree of parallelism between fish and human domestication, particularly, the absence of deliberate selection in both domestication events, and 3) the use of fish as animal models in biomedical research, make fish promising candidate models to identify the epigenetic mechanisms that lead to the emergence of human self-domestication, including their abnormal manifestation in neurodevelopmental disorders.

Comparative epigenomic studies between domesticated animals and humans are expected to demonstrate parallel or contrasting processes operating in addition to traditional genetic aspects [48]. Here, we argue that fish hold great advantages as models to study epigenetic drivers in human self-domestication. To test our argument, we use comparative epigenomic approaches between humans and the European sea bass. The European sea bass was chosen because: 1) 25 years of selective breeding resulted in selective sweeps in genes similar to those found under positive selection in all domesticates tested, e.g., glutamate receptors [49,50], 2) it presents traits of the domestication syndrome shared with those found in terrestrial vertebrates, e.g., depigmentation and cranial changes [35] and 3) epigenetic patterns of domestication have been assessed in four tissues types representative of all three embryonic layers, thus reducing bias due to tissue-specificity [35]. In the present study, we compare epigenetic patterns of domesticated sea bass with epigenetic patterns of 1) AMH as opposed to archaic hominins (Neanderthals and Denisovans), and 2) neurodevelopmental cognitive disorders with an abnormal presentation of traits parallel to the domestication syndrome (SZ, WS and ASD; **Fig. S1**). The goal of these comparisons was to detect genes or pathways consistently altered, or their absence, during the steps of early domestication in European sea bass and humans, with a potential impact on our species-specific distinctive cognition and behaviour.

## 2. Materials and Methods

### 2.1 Data collection

Comparative epigenomic analyses were divided in two major groups including early domesticates of the European sea bass vs 1) AMH and 2) neurodevelopmental cognitive disorders. For this, we compiled five lists of genes identified as differentially methylated in the literature (**Fig. S1)**.

#### 2.1.1 European sea bass early domesticates

In European sea bass, we previously conducted work to generate genome-wide DNA methylation patterns (Reduced Representation Bisulfite Sequencing, RRBS) in fish captured in the wild vs offspring of wild fish reared in hatchery [35]. DNA methylation data from brain, muscle, liver and testis can be accessed through the NCBI Gene Expression Omnibus database with accession codes GSE104366 and GSE125124. Since these data were published, the European sea bass genome has been included in the Ensembl database. The genome assembly v1.0 in Ensembl is the same used for data analysis by [35], however, gene annotation has since been updated according to the Ensembl Gene Annotation pipelines. To facilitate comparative epigenomic analysis with human, we converted the list of genes with differentially methylated regions (DMRs) to the Ensembl genebuild released version from April 2020. To do this, the genomic coordinates (chromosome, start, end position) of DMRs and surrounding 5000 bp regions were intersected with the genebuild Dicentrarchus_labrax.seabass_V1.0.101.gtf. Chromosome names were as in the primary assembly. A total of 1181 unique genes with DMRs were identified in early domesticates.

#### 2.1.2 Anatomically modern human (AMH)

A detailed map of the evolutionary dynamics of DNA methylation in human groups was recently published [30]. DMRs specific to the AMH-lineage as compared to other hominin lineages, i.e. Denisovan and Neanderthal, were identified using a conservative approach to minimize false positives and variability due to factors such as sex or age, as well as DNA methylation data from chimpanzee samples. AMH-lineage DMRs are a set of 873 DMRs that overlap with the gene body or the promoter up to 5000 bp upstream of 588 genes (Supplementary Data 2 of [30]; **Fig. S1**). The list of genes with DMRs was supplied by [30] with UCSC identifiers (IDs) and we used the https://biotools.fr/human/ucsc_id_converter tool to convert them to Ensembl IDs to facilitate comparative epigenomics with the European sea bass.

#### 2.1.3 Neurodevelopmental cognitive disorders

WS has a clear genetic origin with the hemideletion of 28 genes at 7q11.23. Some of these genes, e.g., *BAZ1B*, are involved in epigenetic regulation, such as chromatin remodeling, providing a link to the impact on epigenomic patterns in WS [12]. Differential DNA methylation between patients with WS and healthy individuals as controls has been reported in at least two cases in the literature. DMRs identified using the Infinium HumanMethylation 450 BeadChip array (Illumina) in the blood of 20 WS patients vs 15 healthy controls found DMRs intersecting 551 unique genes [51]. Differentially methylated cytosines (DMCs) were detected more recently in blood of a larger sample of 90 WS patients vs 34 healthy controls using the same array and these intersected with 143 unique genes [52]. The two gene lists were combined for further analysis as genes differentially methylated (DM) in WS, with a total of 624 different genes.

SZ is a complex psychiatric disorder and epigenome-wide association studies (EWAS) have been carried out to explore the role of DNA methylation in SZ pathophysiology, with discordant results. Recently, a metaanalysis of five EWAS datasets was published, including samples taken from different parts of the brain (frontal cortex, cerebellum, hippocampus and prefrontal cortex), between 3 and 47 samples per study and using either the Illumina Infinium Human Methylation 450 Beadchip or Human Methylation 27 BeadChips [53]. A total of 513 genes were commonly DM in combinations of 4-5 EWAS and these were used here for further analysis as the DM genes in SZ.

ASD refers to a group of complex neurodevelopmental disorders with heterogeneous symptoms and underlying etiology. ASD heritability is complex and genetic variants involved are diverse with their number ranging between 1000 and 3000 genes reflecting ASD heterogeneity [54]. Other molecular aspects to better understand ASD include epigenetic variants and several studies were published in the last years. This allowed us to apply more stringent criteria for the inclusion in this study, mainly a minimum number of 15 samples and identification of DMRs which are considered more robust than DMCs only. Genes from four studies published in the last 4 years, thus, included: a) 31 genes with DM and that were at the same time differentially expressed and common in three independent studies based on blood samples [55], b) 181 core genes with DMRs detected using all three approaches in blood cells [54], c) 145 unique differentially expressed genes with DMRs in blood cells from three ASD subphenotypes (severe, intermediate, mild) and a group of combined cases [56] and d) 58 genes with DMRs detected in postmortem brain samples [57]. The four datasets combined led to a list of 411 unique ASD genes.

### 2.2 Comparative analyses

The BioMart data mining tool from Ensembl was used to identify orthologues of human genes from the genome assembly GRCh38.p13 of the European sea bass genome. Duplicate entries were eliminated for further analysis. Thus, we identified unique orthologues as follows: 589 for AMH, 506 for WS, 532 for SZ, and 367 for ASD **(Fig. S1)**. The BioMart tool was used to identify paralogues of the human genes involved in neurodevelopmental cognitive disorders in the human genome (GRCh38.p13), in turn used to identify orthologues in the European sea bass genome. Duplicate entries from the combined list of original orthologues and orthologues of human paralogues were eliminated and the number of homologues finally available for comparative analyses were as follows: 3460 for WS, 4000 for SZ and 2994 for ASD.

Pairwise comparisons were performed with the fish early domesticates (FED) as a reference and one human group as its pair. Thus, 4 pairwise comparison occurred every time: 1) FED vs AMH, 2) FED vs WS, 3) FED vs SZ and 4) FED vs ASD. Overlaps between gene lists were identified and visualized using the InteractiVenn tool [58].

Significance of overlap was tested using Fisher’s exact test for testing the independence of two variables represented by a contingency table. As the genomic background for gene overlap testing, the total number of 23382 genes in the European sea bass genome (Ensembl genebuild released April 2020) was set.

Furthermore, we performed Monte Carlo permutations to test whether overlaps were higher than expected by chance. Random samples of genes were drawn without replacement from the 23883 total gene list according to the specific gene list each time, e.g., to test the overlap of orthologues FED vs AMH, 1181 genes for FED vs 589 genes for AMH were randomly drawn in each iteration. The process was repeated 10000 times and each time the length of the intersection or overlap between the two genes lists was counted. The standard score of permutation was calculated as: observed-mean(permuted)/sd(permuted) and the *p*-value as: times permuted overlap is higher than observed overlap divided by number of permutation (10000). Fisher’s tests and permutations were performed using R (v. 4.0.0) [59] and Rstudio (v. 1.4.1717) [60].

The Enrichr tool was used for enrichment analyses and knowledge discovery of gene sets [61–63]. Enrichment analyses were performed for the initial lists of genes (FED, AMH, WS, SZ and ASD). Enriched pathways from the databases BioPlanet, Wikipathway, Mammalian Phenotype and GO-terms Biological Process were kept for further comparisons which included overlap testing as previously with background the total number of terms found in each library on Erichr. Reduction and visualization of GO-terms was aided by REViGO [64]. IDs of pathways were entered in InteractiVenn to detect overlaps and Fisher’s exact tests were run to detect statistical significance of the overlap. Enrichment analyses were also performed for the genes that overlapped in a pairwise manner between FED genes and homologues (combined lists of orthologues and orthologues of paralogues).

## 3. Results

### 3.1. Differentially-methylated genes during early domestication in fish and in humans are shared

The early stages of domestication are expected to be associated with DNA methylation changes. To compare DNA methylation changes associated with the early stages of domestication between fish and human, two gene lists were retrieved. In FED 1181 genes with DMRs were detected as compared to wild fish. For humans, based on limited availability and accessibility to early AMH domesticate samples, DNA methylation patterns of present-day AMHs compared to other hominins and primates were considered as the most relevant proxy. A total of 589 genes with DMRs were detected as orthologues of AMH. We detected an overlap of 45 genes between FED and AMH and this was significant (Fisher’s test, odds ratio=1.577, *p* = 0.004; **Fig. 1a**). Furthermore, we found 1.7 times more genes in common between the two gene lists than expected by chance alone (z-score=13.62, *p*=3^e-04^; **Fig. 1b**).

**Figure 1.**
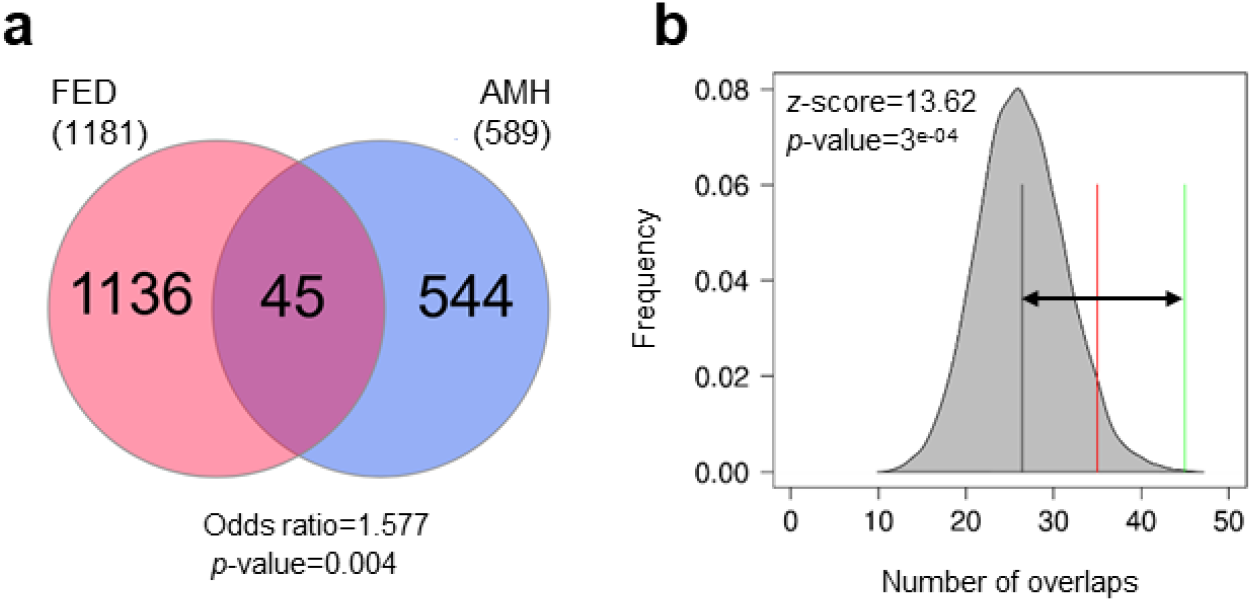
Overlap of genes with epigenetic changes in fish early domesticates (FED) and anatomically modern humans (AMH). The overlap was tested using Fisher’s exact test for count data (a) and permutations (b). The results of permutations are represented as the distribution of number of overlaps (shaded grey area) with mean number of permuted overlaps (black vertical line) and significance threshold set to 0.05 (red line). Observed number of overlaps is shown by the green line and the distance of observed vs expected (random) overlaps is shown with the black arrow. The z-score and the *p*-value indicate the significance of the overlap.

Among the genes with DMRs in both groups (**Table 1**), we detected several genes that were repeatedly found to be involved in domestication in several species. For example, ADAM metallopeptidases with thrombospondin type 1 motifs, ephrin (eph) receptors, members of the integrin family (alpha or beta), or fibroblast growth factor receptors have been detected in other domesticates (see Dataset 1 from [35] for overview and [65– 69] for each species). One of these genes is nuclear factor I X (*NFIX* in humans and *nfxib* in fish) which was found to be in the top 10 genes with DMRs in AMHs showing strong correlation between methylation and expression [30]. Several lines of evidence suggest that hypermethylation of *NFIX* associates with its downregulation in the AMH lineage [30]. In FEDs, *nfixb* was hypermethylated in the testis (+29.98%) but hypomethylated in the muscle tissue (−35.88%). In other tissues, other nuclear factor 1 isoforms contained DMRs: in muscle tissue, nuclear factor 1 a-type contained 2 DMRs with opposite methylation patterns (+20.69% and -42.30%) and in brain tissue, nuclear factor 1 a-type contained 2 hypomethylated DMRs (−27.36% and -34.02) and nuclear factor 1 b-type isoform x2 contained an hypomethylated DMR (−30.14%).

**Table 1.**
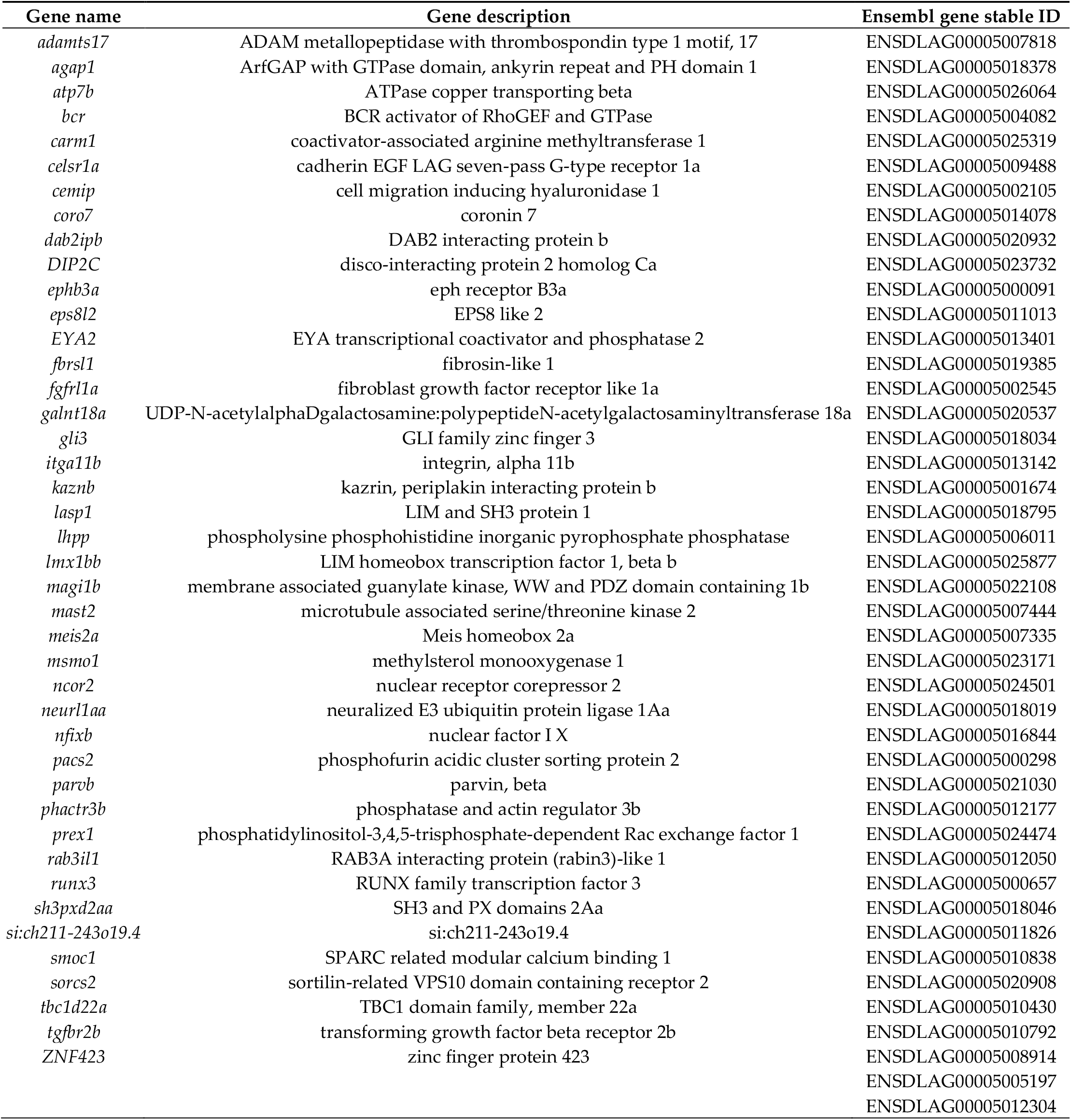
Common genes differentially methylated in fish early domesticates and anatomically modern humans

We performed enrichment analyses to get insight into the functional roles of the overlapping genes. GO Biological Process enrichment analysis highlighted processes such as limb morphogenesis (GO:0035108, *p* = 0.045), histone modifications (GO:0016570, *p* = 0.024), T cell apoptotic processes (GO:0070231, *p*=0.014) or granulocyte activation (GO:0036230, *p* = 0.021) as common (**Fig. 2a**; for full list **Table S1**). Analysis of MGI Mammalian Phenotypes showed enrichment in traits typical of the domestication syndrome, such as abnormal snout morphology (MP:0000443, *p*-adjusted=0.031) or hypopigmentation (MP:0005408, *p*-adjusted=0.034; **Fig. 2b**; for full list **Table S2**). Enrichment of WikiPathways showed that affected pathways include endochondral ossification with skeletal dysplasia (WP4808, *p*=0.008), endochondral ossification (WP474, *p*=0.008) or androgen receptor signaling pathway (WP138, *p*=0.015; for full list **Table S3**).

**Figure 2.**
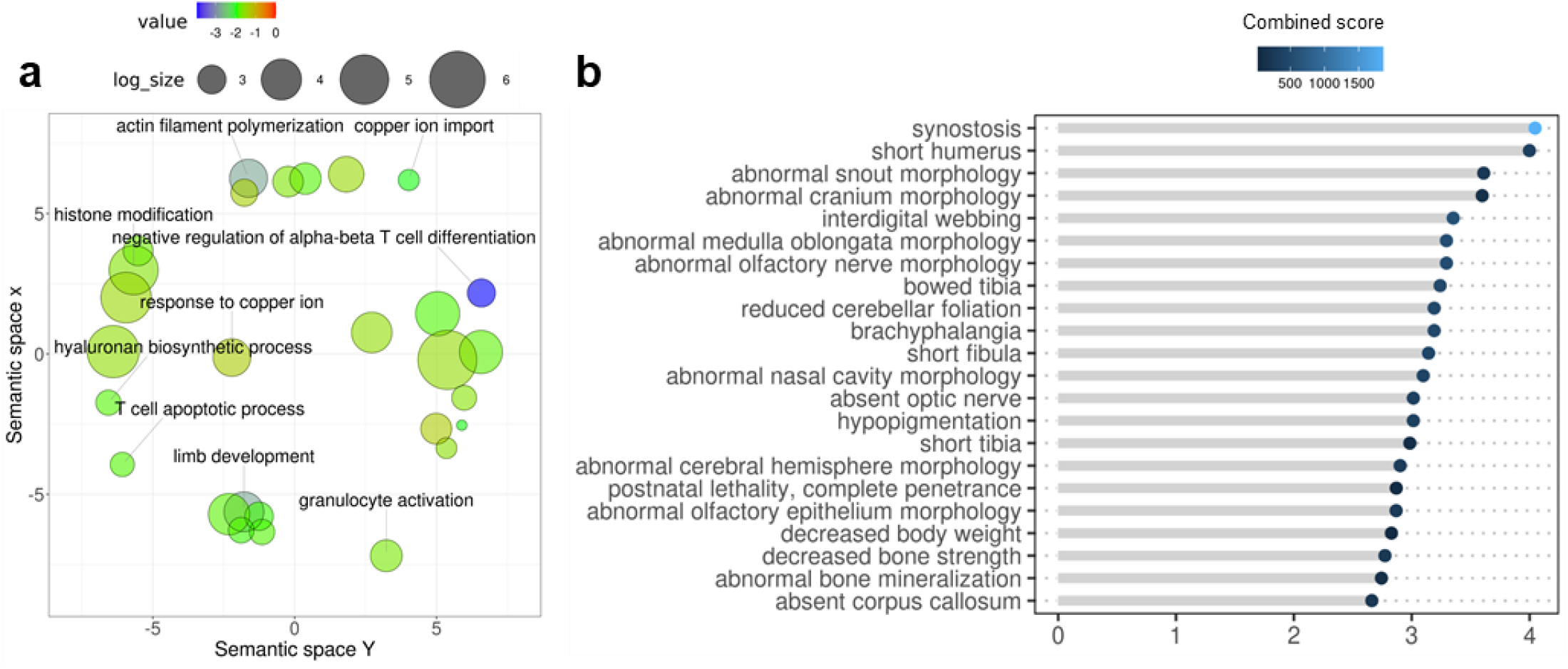
Enrichment analysis of overlapping genes with epigenetic changes in fish early domesticates and anatomically modern humans. a) GO Biological Process terms enrichment where for each GO-term the color indicates the log10-transformed p-value of enrichment. The semantic space x (y-axis) and the semantic space y (x-axis) are the result of multidimensional scaling done by REViGO and represent semantic similarities between GO-terms. b) Pathways of the MGI Mammalian Phenotype 2014 where terms are ranked in descending order according to the -log10-transformed p-value of enrichment and colored according to the combined score estimated by Enrichr.

### 3.2. Early domestication in fish and neurodevelopmental cognitive disorders affect paralogue genes

Genes exhibiting DNA methylation changes in patients with neurodevelopmental cognitive disorders with traits parallel to the domestication syndrome such as SZ, WS and ASD were obtained from the literature. A total of 532 genes with differential methylation (DM) were orthologues to SZ patients, 506 genes with DM in WS patients and 367 genes with DM in ASD patients. These gene lists of orthologues were compared to the genes of FED to evaluate whether DNA methylation in common genes was affected by these conditions. The pairwise overlaps were not significant in all cases, with 28 genes overlapping in SZ (odds ratio=1.04, *p* = 0.439; **Fig. S2a**), 31 overlapping in WS (odds ratio=1.233, *p* = 0.88; **Fig. S2b**) and 23 genes overlapping in ASD (odds ratio=1.262, *p* = 0.169; Fig. **S2c**). Permutation testing for the pairwise comparisons showed that the number of overlaps was within the range expected by chance in the case of SZ (z-score=-0.59, *p*=0.216; **Fig. S2d**) and ASD (z-score=2.61, *p*=0.062; **Fig. S2f**), and only marginally significant in the case of WS (z-score=3.75, *p*=0.049; **Fig. S2e**).

In an attempt to overcome the constraints of the conservative approach applied here for orthologues and since key candidate genes of domestication were present in all pairwise comparisons, e.g. protocadherins, ADAM metallopeptidases, collagens and glutamate receptors, we then focused on comparisons of functional properties. Orthologue genes were submitted for enrichment analyses and pairwise comparisons were performed at the pathway level following the reasoning that similar processes may be affected by different genes. We considered 4 libraries targeted by Enrichr as the most informative in our case: Bioplanet, WikiPathways, GO-terms Biological Process and MGI Mammalian Phenotype. Terms in all 4 libraries were examined for enrichment according to the gene lists we provided (FED, SZ, WS and ASD) and pairwise comparisons of terms were performed as following: 1) FED vs SZ, 2) FED vs WS and 3) FED vs ASD (**Fig. S3**). In 42% of the comparisons, there was no overlap of terms, while in three cases there were between 1 and 4 terms overlapping. The overlaps of terms were significant only in case of SZ for WikiPathways (odds ratio=4.477, *p* = 0.003; **Fig. S3d**) and GO Biological Process (odds ratio=2.442, *p* = 0.002; **Fig. S3g**). WikiPathways included endochondral ossification with skeletal dysplasia (WP4808) and endochondral ossification (WP474) like in the enrichment of orthologue genes overlapping in AMH, but also neural crest differentiation (WP2064). GO Biological Process enriched included development of renal system (GO:0072001), kidney (GO:0001822) or ureteric bud (GO:0001657), as well as regulation of immune cells such as T-helper 17 and alpha-beta T (GO:2000317, GO:0046639 or GO:2000320). Taken together these results indicate that further comparative analyses could reveal more additional similarities.

To investigate the role of gene families, we compared gene lists containing not only the orthologues but also the paralogues of genes. The FED gene list was maintained in the original format and served as the control in the pairwise comparisons completed as above. For the other 3 gene lists (SZ, WS and ASD), paralogues in the human genome were obtained by Biomart, merged with the original genes and then orthologues in the European sea bass genome were identified, resulting in lists containing unique homologues (orthologues and paralogues). The gene lists contained 4000 homologues for SZ, 3460 homologues for WS and 2994 homologues for ASD. Overlap between all pairwise comparisons was significant with 241 genes common in SZ (odds ratio=1.258, *p* = 0.001; **Fig. 3a, Dataset 1**), 236 in WS (odds ratio=1.470, *p* = 4.422^e-07^; **Fig. 3b, Dataset 2**) and 178 overlapping in ASD (odds ratio=1.222, *p* = 0.011; **Fig. 3c, Dataset 3**). Since these gene lists contain ∼8 times more genes than previously, the significance of the overlaps could be attributed to larger numbers. To test whether the number of overlaps could be expected by chance due to large number of genes, we performed Monte Carlo permutations using random sampling of genes from the whole genome as previously. We found that overlaps between gene lists were higher than expected by chance in all cases, including SZ (z-score=49.03, *p*=0; **Fig. 3d**), WS (z-score=33.01, *p*=0; **Fig. 3e**) and ASD (z-score=69.46, *p*=0; **Fig. 3f**). These results confirmed that there were similarities between genes DM early during fish domestication and homologues of genes DM in neurodevelopmental cognitive disorders with domestication syndrome traits.

**Figure 3.**
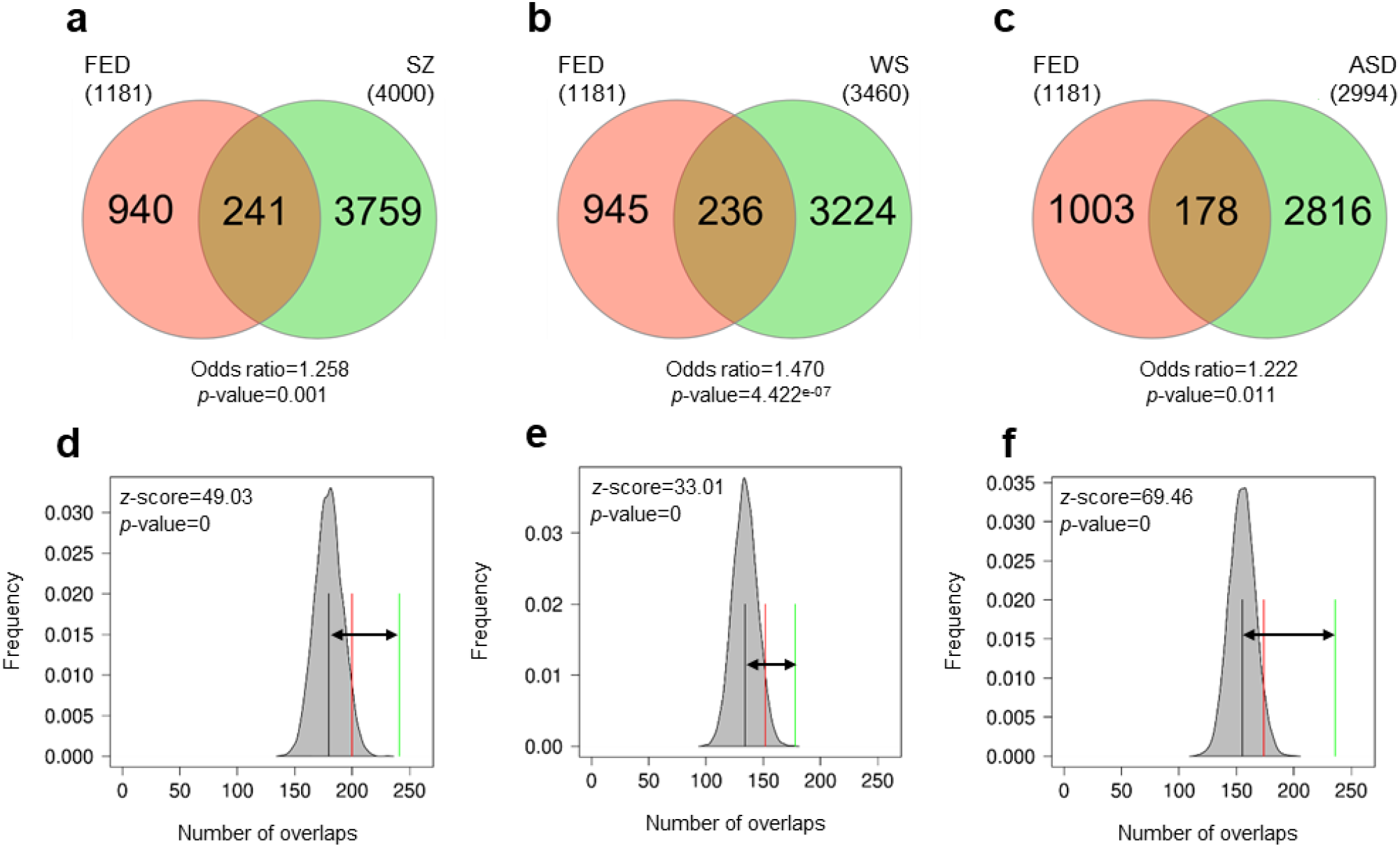
Overlap of homologous genes with epigenetic changes in fish early domesticates (FED) and cognitive disorders. Pairwise comparisons are shown for FED vs schizophrenia (SZ; a,d), Williams syndrome (WS; b,e) and autism spectrum disorders (ASD; c,f). Significance of overlaps were tested using Fisher’s exact test for count data (a-c) and permutations (d-f). The results of permutations are represented as the distribution of number of overlaps (shaded grey areas) with mean number of permuted overlaps (black vertical lines) and significance threshold set to 0.05 (red lines). Observed number of overlaps is indicated by the green lines and the distance of observed vs expected (random) overlaps are shown with the black arrow. The z-scores and the *p*-values indicate the significance of the overlaps.

To evaluate the functional properties of the core overlaps between genes in FED and lists of homologous genes in cognitive disorders, we performed enrichment analysis using Enrichr as previously. Pathways affected in all pairwise comparisons included neural crest differentiation (WP2064), ectoderm differentiation (WP2858), hair follicle development: organogenesis - part 2 of 3 (WP2839), arrhytmogenic right ventricular cardiomyopathy (WP2118; **Fig. 4a-c**, full lists in **Tables S4-6**). Pathways affected in at least two pairwise comparisons included endochondral ossification with skeletal dysplasia (WP4808) like in the core overlap of FED with orthologues of AMH, or also focal adhesion (WP306) and BMP signaling in eyelid development (WP3927) among others (**Fig. 4a-c**).

**Figure 4.**
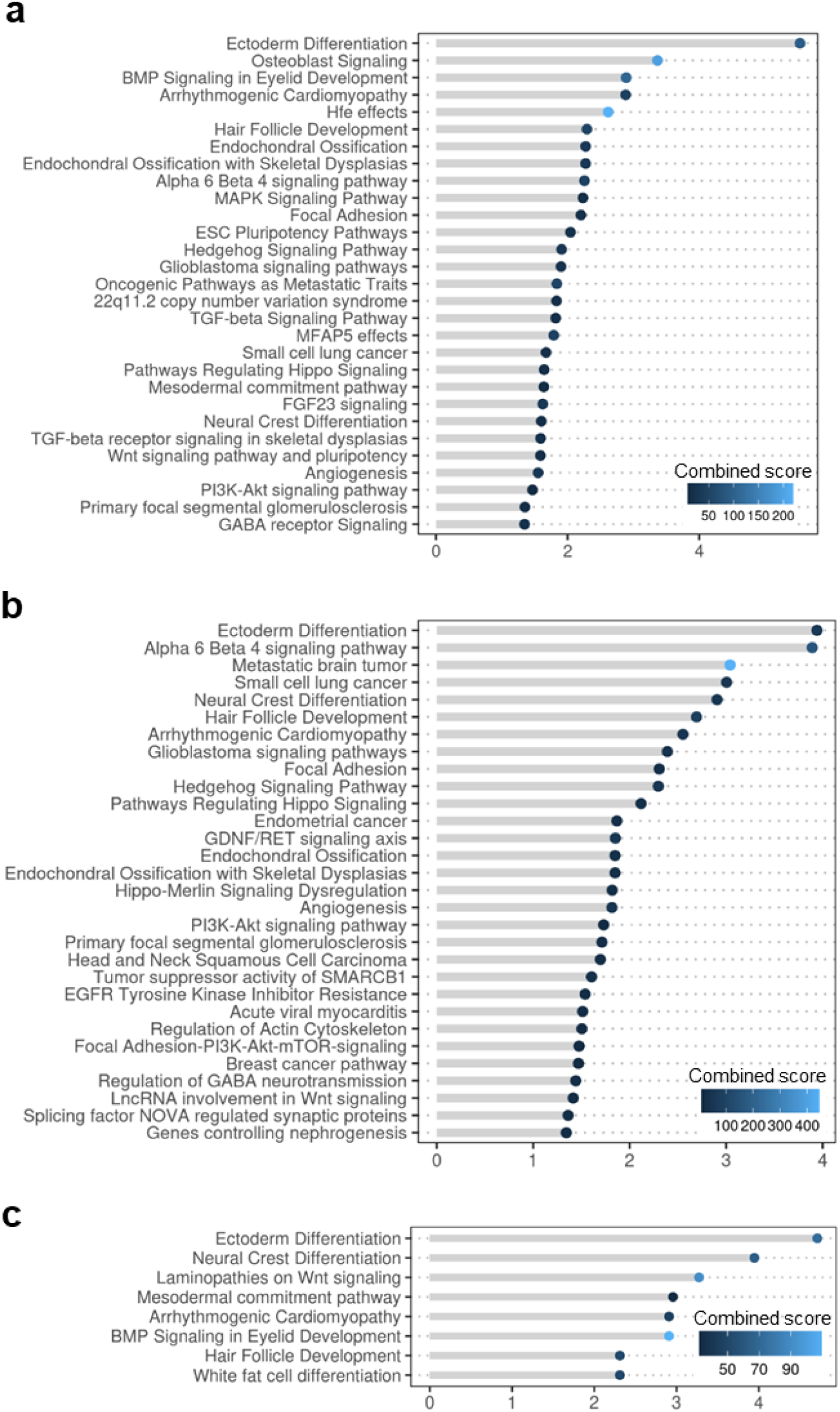
Pathway enrichment of genes with epigenetic changes in fish early domesticates (FED) and homologues of neurodevelopmental cognitive disorders. Pathways of the library Wikipathways enriched in schizophrenia (SZ; a), Williams syndrome (WS; b) and autism spectrum disorders (ASD; c). Terms are ranked in descending order according to the -log10-transformed *p*-value of enrichment and colored according to the combined score estimated by Enrichr.

Further functional analyses included GO-terms of Biological Process. GO-terms affected in all pairwise comparisons included embryonic morphogenesis of skeletal system (GO:0048704), digestive tract (GO:0048557) and organ (GO:0048562), regulation of morphogenesis of a branching structure (GO:0060688), morphogenesis of an epithelium (GO:0002009), neuromuscular junction development (GO:0007528), odontogenesis (GO:0042476) and positive regulation of fibroblast proliferation (GO:0048146; **Fig. 5a-c**; full lists in **Tables S7-9**). In SZ and WS, the extracellular matrix organization was the most significantly enriched GO-term. In ASD, the most significantly enriched GO-term was renal system development and among the enriched GO-terms, we detected glutamatergic synaptic transmission (**Fig. 5c**), a process involving glutamate receptors which have been recognized as affected by domestication across species [35,50].

**Figure 5.**
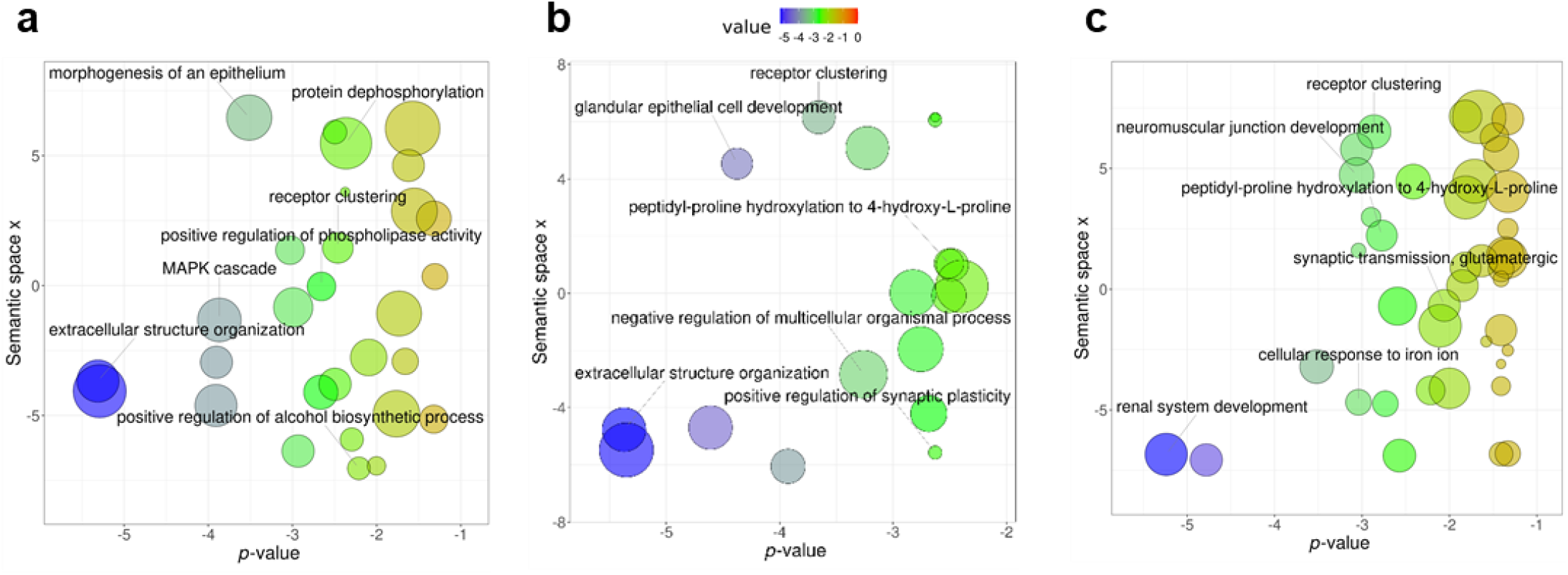
Enrichment of the Gene Ontology (GO) terms of genes with epigenetic changes in fish early domesticates (FED) and homologues of neurodevelopmental cognitive disorders. GO Biological Process terms enrichment in schizophrenia (SZ; a), Williams syndrome (WS; b) and autism spectrum disorders (ASD; c). For each GO-term the color indicates the log10-transformed *p*-value of enrichment which is also represented by the x-axis. The semantic space x (y-axis) is the result of multidimensional scaling done by REViGO and represent semantic similarities between GO-terms.

## 4. Discussion

We have shown that a sizeable portion of epigenetic changes in early fish domesticates occur in similar genes when compared to AMHs, and in similar gene families as in human-specific neurodevelopmental cognitive disorders. Thus, parallel epigenetic changes seem to manifest in independent (self-)domestication processes across vertebrates. Since AMHs exhibit domestication traits and the cognitive disorders studied here (SZ, WS and ASD) exhibit altered phenotypic traits related to the domestication syndrome, all these groups support the hypothesis that humans have been self-domesticated, and that human self-domestication was driven to a great extent by changes in the expression patterns of genes involved in domestication. Our finding that similar genes or gene families exhibited epigenetic changes between human groups and fish provides evidence for domestication as a process affecting similar functional biological properties in vertebrates. Further, it indicates that fish are suitable models for research on epigenomics in human self-domestication, as well as human cognitive disorders.

For the purposes of this study, we compared the lists of genes that exhibited epigenetic changes, measured as differences in DNA methylation. We followed a very conservative approach and included layers of statistical testing, however, some inevitable limitations associated with the nature of the study are present. Genes with epigenetic changes have been pulled from different studies which have used distinct methodologies to interrogate methylation status (e.g., arrays or sequencing) and distinct algorithms to analyze them. However, the data for AMH were deduced from comparisons with reconstructed methylomes using a robust methodology but that dataset lacks methylation data for early AMHs, hence comparisons were performed using data from individuals that were purportedly fully self-domesticated. With regards to neurodevelopmental diseases, due to their often complex etiology, there may be differences in genes detected as DM by different authors who may have used different sampling strategies. Thus, we chose to include only studies which fulfilled stringent criteria. For example, studies involving a very small number of samples, e.g. comparisons of a pair of twins, were excluded. However, we cannot rule out the possibility that the exact gene lists with epigenetic changes may vary slightly when following consistent and unified guidelines for their detection. Furthermore, to detect homologues, the Biomart tool from the Ensembl database was used which is one of the most transparent approaches to perform the task since versions of the genome and annotations can be traced. For the enrichment analyses, genes have to be well characterized and included in the query databases to be informative of the affected pathways. Relying on these bioinformatics resources carries the inherent risks of minor modifications in future updated versions. Nevertheless, the results of this study can be interpreted while taking these limitations into account, since they are based on conservative inclusion criteria and statistical testing and can be used as a step for further research on comparative epigenomics between phylogenetically distant vertebrates.

The human self-domestication hypothesis, as well as the involvement of neural crest cells in human self-domestication, even though attractive, remained mostly supported theoretically until recently. Genomic approaches comparing genes under positive selection between domesticated mammals and AMHs are starting to be used as supportive evidence for the human self-domestication hypothesis [11,70]. Recently, the hypothesis was empirically validated and the role of *BAZ1B*, with an established role in NC induction and migration, was demonstrated [12]. The implication of this gene in morphological and behavioural phenotypes typical of the domestication syndrome via neural crest cell development was further shown using zebrafish as a model [71]. Our comparative results between AMHs and early fish domesticates provide additional support for the role of specific genes in key processes with an impact on (self-)domestication features and suggest a role for epigenetic regulation of their expression. *NFIX* is associated with craniofacial skeletal disease phenotypes and related to speech capabilities, and it has already been highlighted for its role in the development of the AMH face and larynx [12]. Another gene common between early domesticated fish and AMH was GLI family zinc finger 3 (*GLI3*) which is a known transcriptional repressor involved in tissue development, including limb development, and immune system development [72]. *GLI3* has a role during embryogenesis, controlling thalamic development [73], as well as calvarial suture development [74], while in ≈98% of Altaic Neanderthals and Denisovans it contains a mutation that is mildly disruptive [75]. The RUNX family transcription factor 3, *RUNX3*, is involved in the developing spinal cord and also has a role in the language and social regions of brain [76,77]. *SMOC1*, as well as *SMOC2*, play a role in endochondral bone formation and are regulated by another member of the RUNX family transcription factor [78]. *RUNX2* encodes a master transcription factor during vertebrate development involved in the globularization of the human skull/brain. *RUNX2* is also involved in the development of thalamus, which is functionally connected to many genes that are important for brain and language development, and that have experienced changes in our recent evolutionary history [74]. *NCOR2* has already been identified as under selection in dogs [66] and is part of the cranial neural crest gene expression program [79]. The above-mentioned genes participate in the enriched mammalian phenotypes detected which match the domestication syndrome traits, like abnormal cranium morphology, hypopigmentation or decreased body strength, but also in human-distinctive features potentially associated to our self-domestication. Similarly, *GLI3* and *SMOC1* participate in the enriched GO-terms processes related to limb development, including limb morphogenesis and embryonic digit morphogenesis. The GO-term most significantly enriched according to its *p*-value ranking was the negative regulation of alpha-beta T cell differentiation. This is likely to the involvement of the above-mentioned genes, i.e., *RUNX3, GLI3* and *SMOC1*, in the immune system as well. These results together reinforce the role of epigenetics in the regulation of similar genes associated with the domestication syndrome during the early stages of domestication in the absence of deliberate selection, as is the case in both humans and fish. These results also provide support for the view that domestication constitutes an example of “developmental bias”, i.e., when perturbed by an altered environment, complex organisms pursue a limited number of developmental pathways [3].

Neurodevelopmental cognitive disorders in humans have been previously suggested as models for testing the human self-domestication hypothesis [8,15,16]. WS has already been used to gather molecular evidence for the shaping of the human face and behavior underlying self-domestication [12]. Our initial analyses in search of common genes and pathways epigenetically altered in fish domesticates and cognitive disorders was unsuccessful. However, even though orthologue genes seemed to be absent, it was evident that similar gene families were affected, thus, justifying our subsequent approach in the search of paralogues. The lack of common genes could be due to the phylogenetic distance between species and to the nature of conditions tested, i.e., disease phenotypes vs fish under farming conditions.

In SZ and WS, genes of key families were affected including, ADAM metallopeptidases, bone morphogenetic proteins, ephrins, fibroblast growth factors, homeoboxes, laminins and members of the TBC1 domain family. ADAM metallopeptidases and laminins constitute the core members of the most significantly enriched GO-term of overlapping genes in both comparisons: extracellular structure organization. The role of DM genes of the extracellular matrix has already been highlighted in relation to early domestication in fish, and especially for DM changes established already early during development [35]. At the same time, the brain extracellular matrix is known to have multiple roles in brain development and function, and abnormal alteration of this matrix is increasingly acknowledged as a key etiological factor involved in neurological and psychiatric disorders (see [80] for review). In ASD, genes were slightly different and included bone morphogenetic proteins, glutamate receptors, laminins, protocadherins and semaphorins. The migration of neural crest depends on the interaction of receptors, e.g., ephrins and receptors for bone morphogenetic proteins, with extracellular matrix molecules, e.g., laminins and semaphorins [81]. The term neural crest differentiation was enriched in the overlapping groups of genes and consistently found in all three neurodevelopmental disorders, together with ectoderm differentiation, hair follicle development: organogenesis - part 2 of 3 and arrhytmogenic right ventricular cardiomyopathy. Members of this term were *fgfr2, pax3, axin2, hdac10, cdh2, hes1, tfap2a, tfap2b* and *tcf7l1*. Disorders of the processes related to the neural crest are often regarded as underlying SZ, WS and ASD [81]. FGF has an essential embryonic function during vertebrate development and Fgf signaling and has been shown to serve as a target for selection during domestication [82]. In ASD, paralogues of two key genes found in the AMH comparison were also identified as epigenetically altered, i.e., *runx3* and *gli3*. This reinforces the idea that parallel processes are involved in self-domesticated phenotype emergence, either evolutionary or pathologically, supporting the view that cognitive diseases can result from changes in genes involved in human evolution [83,84]. Together these results show that epigenetic changes occur in similar gene families in independent models of early (self-)domestication and that several of these genes have already an established role in the neural crest and other processes recognized as affected by (self-)domestication.

Fish as animal models have long been used in basic science. Small teleost fish, like zebrafish or medaka, have been recently considered as models to study human neurological disorders including ASD [85], peripheral neuropathy [86] and for behavioral neuroscience [87] since they possess several key advantages [88]. First, they consist of a phylogenetically diverse group with species that have evolved phenotypes naturally mimicking human diseases, called “evolutionary mutant models”[89–91]. Cross-species comparisons allow to identify the best models to study a specific physiological pathway [39]. Furthermore, in model species like zebrafish, genetic mutants for specific genes can be easily generated. Second, since they are vertebrates, their brain basic structure and function exhibit similarities to humans showing conserved neuronal circuitry [92]. Third, teleost genomes show homology with 70% of genes associated to human diseases [93,94]. Fourth, model fish species larvae are transparent, offering the opportunity for direct observation of the central nervous system during development [95]. Thus, the use of fish models to study neurodevelopmental cognitive disorders exhibiting (self-)domestication-related features has already a sound basis on previous research. Indeed, zebrafish has been used as a model for the three disorders studied here, SCZ [96], WS [97] and ASD [98]. Our findings that homologue genes were differentially methylated in both human disorders and early fish domesticates provides further evidence for the use of fish as models to study the epigenomic regulation implicated in self-domestication-related human phenotypes, which has proven to be key source of the human uniqueness [5].

For research related to the human self-domestication hypothesis, fish not only possess the above-mentioned advantages, but also show a key similarity distinct from most farm animals: fish domestication and human self-domestication took place in absence of deliberate selection. Our result that DNA methylation changes in fish early domesticates and human groups manifested in overlapping genes supports the implication of epigenetic mechanisms in domestication as a process of adaptation to a human-made environment, but also in the generation of such human-made environments, at least, the environment resulting from our self-domestication. A recent study used zebrafish investigated the role of neural crest in the morphological and behavioral domesticated phenotypes in human self-domestication [71]. They found that a loss of function of the key gene in WS and for the neural crest, *baz1b*, identified as important previously in humans as well [12], resulted in mild neural crest deficiencies during development and behavioral changes related to stress and sociality in adulthood [71]. Furthermore, comparative genomics using domesticated mammals have already been used to shed light to the human self-domestication hypothesis [11]. Together these results show that fish can be implemented in comparative (epi)genomics approaches and functional studies to test the human self-domestication hypothesis.

## 5. Conclusions

We have demonstrated the occurrence of parallel epigenetic changes during independent domestication events in phylogenetically distant vertebrates. These events were driven by living in human-made environments, including the creation of the very human-specific niche through self-domestication, rather than by intentional selection. Epigenetic changes could be the first level of response to a new environment that could later be genomically integrated. An important part of these parallel epigenetic changes arises in genes associated with the neural crest, further supporting the involvement of mild deficits during neural crest development in the emergence of the domestication syndrome. Other common epigenetic changes manifest in genes with neurological or morphological functions that have been associated with the domestication phenotype, including human self-domestication. These findings contribute to our understanding of the initial molecular changes happening during early (self-)domestication and pave the way for future studies using fish as models to investigate epigenetic changes as drivers of human-self domestication, but also as etiological factors of human-specific cognitive diseases.

## Supplementary Materials

The following supporting information can be downloaded at: www.mdpi.com/xxx/s1, Figure S1: Conceptual design of the study; Figure S2: Overlap of orthologue genes differentially methylated in fish early domesticates (FED) and cognitive disorders; Figure S3: Overlap of pathways enriched associated with orthologue genes differentially methylated in fish early domesticates (FED) and cognitive disorders; Table S1: Enrichment of Gene Ontology (GO) Biological Process (BP) terms associated with genes shared between early fish domesticates and anatomically modern humans; Table S2: Enrichment of Mammalian Phenotype (2014) terms associated with genes shared between early fish domesticates and anatomically modern humans; Table S3: Enrichment of Wikipathways associated with genes shared between early fish domesticates and human groups with schizophrenia; Table S4: Enrichment of Wikipathways associated with genes shared between early fish domesticates and human groups with Williams syndrome; Table S5: Enrichment of Wikipathways associated with genes shared between early fish domesticates and human groups with autism spectrum disorders; Table S6: Enrichment of Gene Ontology (GO) Biological Process (BP) terms associated with genes shared between early fish domesticates and human groups with schizophrenia; Table S7: Enrichment of Gene Ontology (GO) Biological Process (BP) terms associated with genes shared between early fish domesticates and human groups with Williams syndrome; Table S8: Enrichment of Gene Ontology (GO) Biological Process (BP) terms associated with genes shared between early fish domesticates and human groups with autism spectrum disorders; Dataset 1: homologue genes with epigenetic changes in human groups with schrizophrenia; Dataset 2: homologue genes with epigenetic changes in human groups with Williams syndrome; Dataset 3: homologue genes with epigenetic changes in human groups with autism spectrum disorders.

## Author Contributions

Conceptualization, D.A., F.P., M.W., A.B.B.; methodology, D.A.; formal analysis, D.A.; investigation, D.A.; data curation, D.A.; writing—original draft preparation, D.A.; writing—review and editing, F.P., M.W., A.B.B.; visualization, D.A.; supervision, F.P., M.W., A.B.B. All authors have read and agreed to the published version of the manuscript.

## Funding

DA was supported by a Marsden grant Te Pūtea Rangahau managed by the Royal Society of New Zealand Te Apārangi to MW (MFP-PAF-1801). This research was supported in part by grant PID2020-114516GB-I00 funded by MCIN/AEI/ 10.13039/501100011033 to ABB, and AEI grant PID2019-108888RB-I00 to FP.

## Data Availability Statement

Data used in this study have been previously published and the details are included in the Materials and Methods section. Any new data generated from re-analysis are included as Supplementary Materials.

## Acknowledgments

Author FP acknowledges the ‘Severo Ochoa Centre of Excellence’ accreditation (CEX2019-000928-S).

## Conflicts of Interest

The authors declare no conflict of interest.

## Notes

### Competing Interest Statement

The authors have declared no competing interest.

