## Supplementary Information for "Fish as model systems to study epigenetic drivers in human self-domestication and neurodevelopmental cognitive disorders"

**neurodevelopmental cognitive disorders**

Dafni Anastasiadi, Francesc Piferrer, Maren Wellenreuther and Antonio Benítez-Burraco

**Contents:**

Figures S1 to S3

Tables S1 to S9

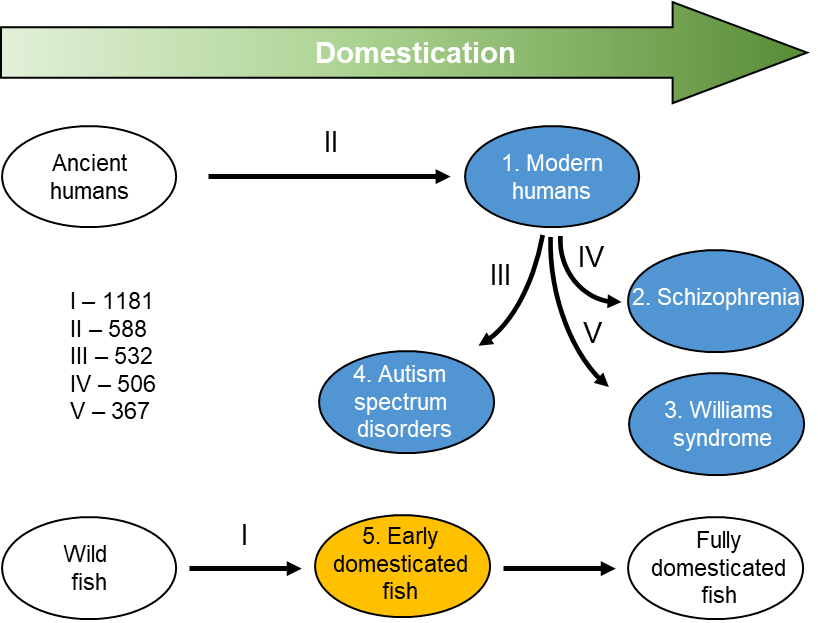

**Figure S1**. Conceptual design of the study. The level of domestication is shown from left to right, from less domesticated phenotypes (ancient humans and wild fish) to more domesticated (patients with schizophrenia or Williams syndrome and fish selected for specific phenotypes). The five groups used in our study are shown by their colored background (blue, humans; orange, fish). The number of genes showing epigenetic changes between groups are shown on the left after the latin numbers.

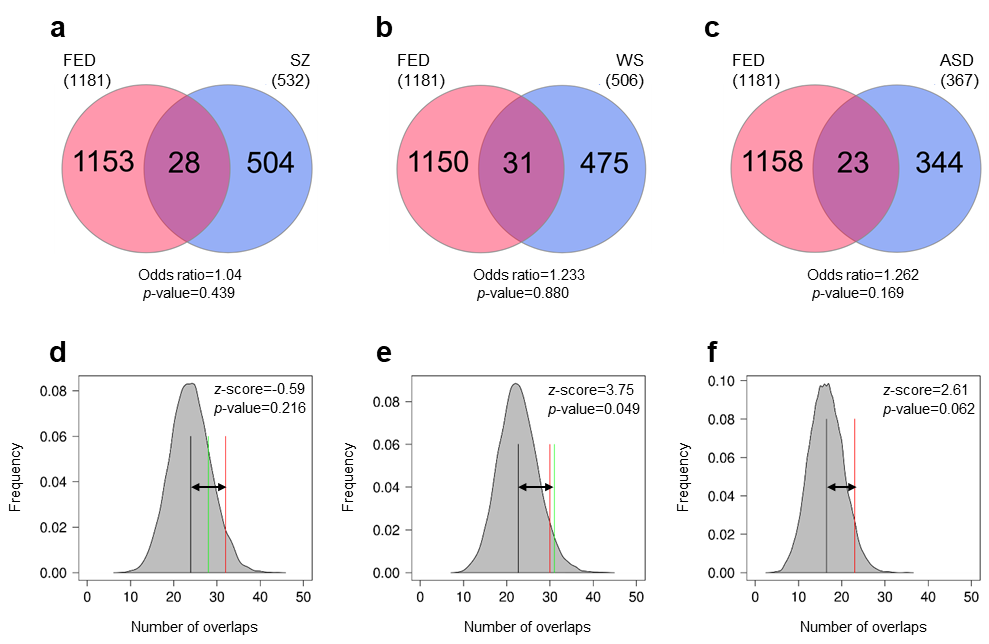

**Figure S2.** Overlap of orthologue genes with epigenetic changes in fish early domesticates (FED) and neurodevelopmental cognitive disorders. Pairwise comparisons are shown for FED vs schizophrenia (SZ; a,d), Williams syndrome (WS; b,e) and autism spectrum disorders (ASD; c,f). The overlaps were tested using Fisher’s exact test for count data (a-c) and permutations (d-f). The results of permutations are represented as the distribution of number of overlaps (shaded grey areas) with mean number of permuted overlaps (black vertical lines) and significance threshold set to 0.05 (red lines). Observed number of overlaps is shown by the green lines and the distance of observed vs expected (random) overlaps is shown with the black arrow. The z-scores and the p-values indicate the significance of the overlaps.

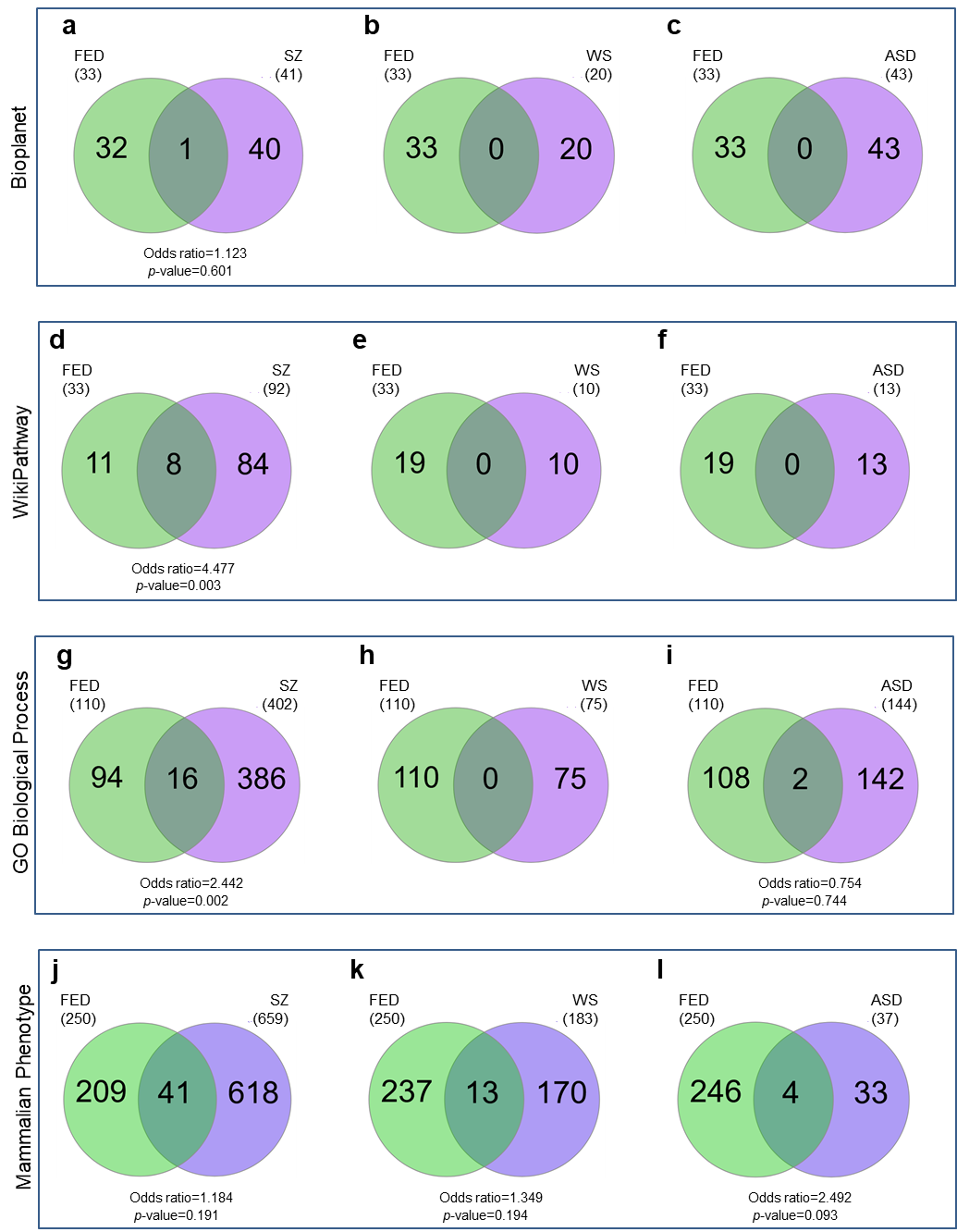

**Figure S3**. Overlap of pathways enriched associated with orthologue genes with epigenetic changes in fish early domesticates (FED) and cognitive disorders. Pairwise comparisons are shown for FED vs schizophrenia (SZ; a, d, g, j), Williams syndrome (WS; b, e, h, k) and autism spectrum disorders (ASD; c, f, I, l). Enriched terms from Bioplanet (a-c), WikiPathways (d-f), Gene Ontology (GO) Biological Process (g-i) and Mammalian Phenotype (j-l) were detected using Enrichr.

**Table S1**. Enrichment of Gene Ontology (GO) Biological Process (BP) terms associated with genes shared between early fish domesticates and anatomically modern humans

| **Biological process GO** | **ID** | **Overlap** | **P-value** | **Adjusted P-value** | **Odds Ratio** | **Combined Score** | **Genes** |
| --- | --- | --- | --- | --- | --- | --- | --- |
| negative regulation of alpha-beta T cell differentiation | GO:0046639 | 2/8 | 1.20E-04 | 0.028 | 166.3 | 1501.6 | *RUNX3;GLI3* |
| positive regulation of alpha-beta T cell differentiation | GO:0046638 | 2/14 | 3.86E-04 | 0.045 | 83.1 | 653.3 | *RUNX3;GLI3* |
| actin filament polymerization | GO:0030041 | 2/29 | 0.002 | 0.098 | 36.9 | 235.7 | *PREX1;CORO7* |
| limb development | GO:0060173 | 2/29 | 0.002 | 0.098 | 36.9 | 235.7 | *SMOC1;GLI3* |
| actin polymerization or depolymerization | GO:0008154 | 2/50 | 0.005 | 0.150 | 20.7 | 110.1 | *PREX1;CORO7* |
| protein polymerization | GO:0051258 | 2/59 | 0.007 | 0.150 | 17.5 | 87.1 | *PREX1;CORO7* |
| regulation of Rho protein signal transduction | GO:0035023 | 2/73 | 0.010 | 0.150 | 14.0 | 64.1 | *BCR;EPS8L2* |
| negative regulation of CD4-positive, alpha-beta T cell differentiation | GO:0043371 | 1/5 | 0.010 | 0.150 | 121.7 | 554.9 | *RUNX3* |
| positive regulation of CD8-positive, alpha-beta T cell differentiation | GO:0043378 | 1/5 | 0.010 | 0.150 | 121.7 | 554.9 | *RUNX3* |
| positive regulation of protein kinase C activity | GO:1900020 | 1/5 | 0.010 | 0.150 | 121.7 | 554.9 | *CEMIP* |
| copper ion import | GO:0015677 | 1/5 | 0.010 | 0.150 | 121.7 | 554.9 | *ATP7B* |
| peripheral nervous system neuron differentiation | GO:0048934 | 1/5 | 0.010 | 0.150 | 121.7 | 554.9 | *RUNX3* |
| regulation of protein kinase C activity | GO:1900019 | 1/5 | 0.010 | 0.150 | 121.7 | 554.9 | *CEMIP* |
| mesodermal cell fate commitment | GO:0001710 | 1/6 | 0.013 | 0.150 | 97.3 | 426.2 | *EYA2* |
| peripheral nervous system neuron development | GO:0048935 | 1/6 | 0.013 | 0.150 | 97.3 | 426.2 | *RUNX3* |
| regulation of Ras protein signal transduction | GO:0046578 | 2/86 | 0.014 | 0.150 | 11.8 | 50.4 | *BCR;EPS8L2* |
| T cell apoptotic process | GO:0070231 | 1/7 | 0.015 | 0.150 | 81.1 | 342.8 | *GLI3* |
| nose development | GO:0043584 | 1/7 | 0.015 | 0.150 | 81.1 | 342.8 | *GLI3* |
| regulation of CD8-positive, alpha-beta T cell differentiation | GO:0043376 | 1/7 | 0.015 | 0.150 | 81.1 | 342.8 | *RUNX3* |
| mitochondrion-endoplasmic reticulum membrane tethering | GO:1990456 | 1/7 | 0.015 | 0.150 | 81.1 | 342.8 | *PACS2* |
| hyaluronan biosynthetic process | GO:0030213 | 1/7 | 0.015 | 0.150 | 81.1 | 342.8 | *CEMIP* |
| sensory perception of mechanical stimulus | GO:0050954 | 2/88 | 0.015 | 0.150 | 11.6 | 48.8 | *CEMIP;EPS8L2* |
| sensory perception of sound | GO:0007605 | 2/91 | 0.016 | 0.150 | 11.2 | 46.4 | *CEMIP;EPS8L2* |
| regulation of lipid metabolic process | GO:0019216 | 2/92 | 0.016 | 0.150 | 11.0 | 45.6 | *NCOR2;CARM1* |
| regulation of alpha-beta T cell differentiation | GO:0046637 | 1/8 | 0.017 | 0.150 | 69.5 | 284.6 | *GLI3* |
| positive regulation of CD8-positive, alpha-beta T cell activation | GO:2001187 | 1/9 | 0.019 | 0.150 | 60.8 | 241.9 | *RUNX3* |
| histone arginine methylation | GO:0034969 | 1/9 | 0.019 | 0.150 | 60.8 | 241.9 | *CARM1* |
| negative regulation of alpha-beta T cell activation | GO:0046636 | 1/9 | 0.019 | 0.150 | 60.8 | 241.9 | *GLI3* |
| sensory organ morphogenesis | GO:0090596 | 1/9 | 0.019 | 0.150 | 60.8 | 241.9 | *GLI3* |
| activation of GTPase activity | GO:0090630 | 2/105 | 0.021 | 0.152 | 9.6 | 37.5 | *BCR;TBC1D22A* |
| granulocyte activation | GO:0036230 | 1/10 | 0.021 | 0.152 | 54.1 | 209.4 | *PREX1* |
| protein localization to phagophore assembly site | GO:0034497 | 1/11 | 0.023 | 0.152 | 48.7 | 183.8 | *PACS2* |
| copper ion transport | GO:0006825 | 1/11 | 0.023 | 0.152 | 48.7 | 183.8 | *ATP7B* |
| histone modification | GO:0016570 | 2/114 | 0.024 | 0.152 | 8.9 | 33.1 | *EYA2;CARM1* |
| purine nucleoside monophosphate biosynthetic process | GO:0009127 | 1/12 | 0.025 | 0.152 | 44.2 | 163.3 | *LHPP* |
| negative regulation of gene silencing by miRNA | GO:0060965 | 1/12 | 0.025 | 0.152 | 44.2 | 163.3 | *NCOR2* |
| regulation of CD4-positive, alpha-beta T cell differentiation | GO:0043370 | 1/12 | 0.025 | 0.152 | 44.2 | 163.3 | *RUNX3* |
| negative regulation of production of miRNAs involved in gene silencing by miRNA | GO:1903799 | 1/12 | 0.025 | 0.152 | 44.2 | 163.3 | *NCOR2* |
| negative regulation of CD4-positive, alpha-beta T cell activation | GO:2000515 | 1/13 | 0.027 | 0.153 | 40.5 | 146.5 | *RUNX3* |
| positive regulation of ruffle assembly | GO:1900029 | 1/13 | 0.027 | 0.153 | 40.5 | 146.5 | *EPS8L2* |
| negative regulation of T cell differentiation | GO:0045581 | 1/14 | 0.029 | 0.153 | 37.4 | 132.5 | *GLI3* |
| T cell differentiation in thymus | GO:0033077 | 1/14 | 0.029 | 0.153 | 37.4 | 132.5 | *GLI3* |
| ribonucleoside monophosphate biosynthetic process | GO:0009156 | 1/14 | 0.029 | 0.153 | 37.4 | 132.5 | *LHPP* |
| cellular copper ion homeostasis | GO:0006878 | 1/14 | 0.029 | 0.153 | 37.4 | 132.5 | *ATP7B* |
| regulation of primary metabolic process | GO:0080090 | 2/130 | 0.030 | 0.157 | 7.7 | 27.0 | *NCOR2;CARM1* |
| negative regulation of androgen receptor signaling pathway | GO:0060766 | 1/15 | 0.031 | 0.157 | 34.7 | 120.6 | *NCOR2* |
| purine ribonucleoside monophosphate biosynthetic process | GO:0009168 | 1/16 | 0.033 | 0.157 | 32.4 | 110.5 | *LHPP* |
| neutrophil activation | GO:0042119 | 1/16 | 0.033 | 0.157 | 32.4 | 110.5 | *PREX1* |
| Golgi to endosome transport | GO:0006895 | 1/16 | 0.033 | 0.157 | 32.4 | 110.5 | *CORO7* |
| protein dephosphorylation | GO:0006470 | 2/139 | 0.034 | 0.157 | 7.2 | 24.4 | *EYA2;LHPP* |
| copper ion homeostasis | GO:0055070 | 1/17 | 0.035 | 0.157 | 30.4 | 101.8 | *ATP7B* |
| regulation of small GTPase mediated signal transduction | GO:0051056 | 2/141 | 0.035 | 0.157 | 7.1 | 23.8 | *BCR;PREX1* |
| hyaluronan catabolic process | GO:0030214 | 1/18 | 0.037 | 0.157 | 28.6 | 94.2 | *CEMIP* |
| embryonic digit morphogenesis | GO:0042733 | 1/18 | 0.037 | 0.157 | 28.6 | 94.2 | *GLI3* |
| regulation of production of miRNAs involved in gene silencing by miRNA | GO:1903798 | 1/18 | 0.037 | 0.157 | 28.6 | 94.2 | *NCOR2* |
| negative regulation of RNA metabolic process | GO:0051253 | 1/19 | 0.039 | 0.159 | 27.0 | 87.5 | *NCOR2* |
| positive regulation of alpha-beta T cell activation | GO:0046635 | 1/20 | 0.041 | 0.159 | 25.6 | 81.6 | *GLI3* |
| purine ribonucleoside monophosphate metabolic process | GO:0009167 | 1/20 | 0.041 | 0.159 | 25.6 | 81.6 | *LHPP* |
| embryonic digestive tract development | GO:0048566 | 1/20 | 0.041 | 0.159 | 25.6 | 81.6 | *GLI3* |
| endoplasmic reticulum calcium ion homeostasis | GO:0032469 | 1/20 | 0.041 | 0.159 | 25.6 | 81.6 | *PACS2* |
| regulation of cell differentiation | GO:0045595 | 2/156 | 0.042 | 0.159 | 6.4 | 20.3 | *SMOC1;RUNX3* |
| focal adhesion assembly | GO:0048041 | 1/21 | 0.043 | 0.159 | 24.3 | 76.4 | *BCR* |
| limb morphogenesis | GO:0035108 | 1/22 | 0.045 | 0.159 | 23.2 | 71.7 | *GLI3* |
| cell projection assembly | GO:0030031 | 1/22 | 0.045 | 0.159 | 23.2 | 71.7 | *PARVB* |
| negative regulation of signal transduction in absence of ligand | GO:1901099 | 1/23 | 0.047 | 0.159 | 22.1 | 67.5 | *EYA2* |
| negative regulation of smoothened signaling pathway | GO:0045879 | 1/23 | 0.047 | 0.159 | 22.1 | 67.5 | *GLI3* |
| negative regulation of extrinsic apoptotic signaling pathway in absence of ligand | GO:2001240 | 1/23 | 0.047 | 0.159 | 22.1 | 67.5 | *EYA2* |
| response to copper ion | GO:0046688 | 1/23 | 0.047 | 0.159 | 22.1 | 67.5 | *ATP7B* |
| hyaluronan metabolic process | GO:0030212 | 1/23 | 0.047 | 0.159 | 22.1 | 67.5 | *CEMIP* |
| regulation of ruffle assembly | GO:1900027 | 1/24 | 0.049 | 0.161 | 21.1 | 63.7 | *EPS8L2* |
| positive regulation of protein targeting to membrane | GO:0090314 | 1/24 | 0.049 | 0.161 | 21.1 | 63.7 | *CEMIP* |

Note. The full enrichment analysis for genes shared between fish early domesticates and anatomically modern human can be accessed here: <https://maayanlab.cloud/Enrichr/enrich?dataset=5d2c407bedd97bc283c4828e2b14c47e>

**Table S2**. Enrichment of Mammalian Phenotype (2014) terms associated with genes shared between early fish domesticates and anatomically modern humans

| **Mammalian Phenotype** | **ID** | **Overlap** | ***p*-value** | **Adjusted *p*-value** | **Odds Ratio** | **Combined Score** | **Genes** |
| --- | --- | --- | --- | --- | --- | --- | --- |
| synostosis | MP:0000566 | 2/7 | 8.98E-05 | 0.025 | 199.5 | 1859.2 | *SMOC1;GLI3* |
| short humerus | MP:0004351 | 3/43 | 1.00E-04 | 0.025 | 38.3 | 352.7 | *ADAMTS17;RUNX3;GLI3* |
| abnormal snout morphology | MP:0000443 | 3/58 | 2.45E-04 | 0.031 | 27.8 | 231.4 | *SMOC1;AGAP1;EPS8L2* |
| abnormal cranium morphology | MP:0000438 | 4/147 | 2.52E-04 | 0.031 | 14.6 | 120.8 | *SMOC1;AGAP1;RUNX3;GLI3* |
| interdigital webbing | MP:0000571 | 2/15 | 4.44E-04 | 0.032 | 76.7 | 592.1 | *SMOC1;GLI3* |
| abnormal olfactory nerve morphology | MP:0005236 | 2/16 | 5.07E-04 | 0.032 | 71.2 | 540.4 | *ZNF423;GLI3* |
| abnormal medulla oblongata morphology | MP:0000846 | 2/16 | 5.07E-04 | 0.032 | 71.2 | 540.4 | *ZNF423;GLI3* |
| bowed tibia | MP:0004358 | 2/17 | 5.74E-04 | 0.032 | 66.5 | 496.1 | *SMOC1;GLI3* |
| brachyphalangia | MP:0002543 | 2/18 | 6.45E-04 | 0.032 | 62.3 | 457.8 | *ADAMTS17;GLI3* |
| reduced cerebellar foliation | MP:0009719 | 2/18 | 6.45E-04 | 0.032 | 62.3 | 457.8 | *ZNF423;GLI3* |
| short fibula | MP:0002765 | 2/19 | 7.20E-04 | 0.032 | 58.7 | 424.4 | *SMOC1;GLI3* |
| abnormal nasal cavity morphology | MP:0002237 | 2/20 | 7.99E-04 | 0.033 | 55.4 | 395.1 | *ZNF423;GLI3* |
| hypopigmentation | MP:0005408 | 2/22 | 9.68E-04 | 0.034 | 49.8 | 345.9 | *ATP7B;GLI3* |
| absent optic nerve | MP:0001333 | 2/22 | 9.68E-04 | 0.034 | 49.8 | 345.9 | *SMOC1;GLI3* |
| short tibia | MP:0002764 | 4/214 | 0.00104 | 0.034 | 9.9 | 68.0 | *PACS2;SMOC1;ADAMTS17;GLI3* |
| abnormal cerebral hemisphere morphology | MP:0008540 | 2/25 | 0.00125 | 0.037 | 43.3 | 289.6 | *ATP7B;GLI3* |
| postnatal lethality, complete penetrance | MP:0011085 | 5/394 | 0.00135 | 0.037 | 6.8 | 44.9 | *ATP7B;SMOC1;ZNF423;RUNX3;GLI3* |
| abnormal olfactory epithelium morphology | MP:0008789 | 2/26 | 0.00136 | 0.037 | 41.5 | 274.3 | *ZNF423;GLI3* |
| decreased body weight | MP:0001262 | 9/1329 | 0.00149 | 0.039 | 3.9 | 25.1 | *NCOR2;PREX1;LASP1;ATP7B;SMOC;* |
| decreased bone strength | MP:0004991 | 2/29 | 0.00169 | 0.042 | 36.9 | 235.7 | *ADAMTS17;PARVB;ZNF423;RUNX3* |
| abnormal bone mineralization | MP:0002896 | 3/115 | 0.00181 | 0.042 | 13.6 | 86.1 | *NCOR2;RUNX3* |
| absent corpus callosum | MP:0002196 | 2/33 | 0.00218 | 0.049 | 32.1 | 197.0 | *PARVB;RUNX3;GLI3* |

Note. The full enrichment analysis for genes shared between fish early domesticates and anatomically modern human can be accessed here: <https://maayanlab.cloud/Enrichr/enrich?dataset=5d2c407bedd97bc283c4828e2b14c47e>

**Table S3**. Enrichment of Wikipathways associated with genes shared between early fish domesticates and anatomically modern humans

| **Wikipathways** | **Overlap** | **P-value** | **Adjusted P-value** | **Odds Ratio** | **Combined Score** | **Genes** |
| --- | --- | --- | --- | --- | --- | --- |
| TGF-beta Receptor Signaling | 2/54 | 5.75E-03 | 0.066 | 19.14 | 98.74 | *ZNF423;RUNX3* |
| TGF-beta receptor signaling in skeletal dysplasias | 2/58 | 6.61E-03 | 0.066 | 17.77 | 89.20 | *ZNF423;RUNX3* |
| Endochondral Ossification with Skeletal Dysplasias | 2/64 | 7.99E-03 | 0.066 | 16.05 | 77.49 | *RUNX3;GLI3* |
| Endochondral Ossification | 2/64 | 0.008 | 0.066 | 16.05 | 77.49 | *RUNX3;GLI3* |
| Androgen receptor signaling pathway | 2/90 | 0.015 | 0.101 | 11.29 | 47.16 | *NCOR2;CARM1* |
| Tgif disruption of Shh signaling | 1/9 | 0.019 | 0.103 | 60.82 | 241.88 | *GLI3* |
| NO metabolism in cystic fibrosis | 1/13 | 0.027 | 0.121 | 40.54 | 146.48 | *CARM1* |
| Cholesterol Biosynthesis Pathway | 1/15 | 0.031 | 0.121 | 34.75 | 120.64 | *MSMO1* |
| Hedgehog Signaling Pathway Netpath | 1/16 | 3.31E-02 | 0.121 | 32.43 | 110.53 | *GLI3* |
| Imatinib and Chronic Myeloid Leukemia | 1/20 | 4.12E-02 | 0.136 | 25.60 | 81.64 | *BCR* |
| GDNF/RET signaling axis | 1/23 | 4.72E-02 | 0.141 | 22.10 | 67.47 | *GLI3* |

Note. The full enrichment analysis for genes shared between fish early domesticates and anatomically modern human can be accessed here: <https://maayanlab.cloud/Enrichr/enrich?dataset=5d2c407bedd97bc283c4828e2b14c47e>

**Table S4**. Enrichment of Wikipathways associated with genes shared between early fish domesticates and human groups with schizophrenia

| **Wikipathways** | **Overlap** | **P-value** | **Adjusted P-value** | **Odds Ratio** | **Combined Score** | **Genes** |
| --- | --- | --- | --- | --- | --- | --- |
| GABA receptor Signaling | 2/31 | 0.045 | 0.275 | 6.3 | 19.5 | *SLC32A1;GABRD* |
| Hair Follicle Development | 3/32 | 0.005 | 0.101 | 9.5 | 50.0 | *LAMA5;PDGFRA;FGFR2* |
| Alpha 6 Beta 4 signaling pathway | 3/33 | 0.006 | 0.101 | 9.1 | 47.5 | *LAMA5;LAMA2;LAMA3* |
| Hedgehog Signaling Pathway | 3/44 | 0.012 | 0.158 | 6.7 | 29.4 | *SMURF2;BOC;FBXL17* |
| TGF-beta receptor signaling in skeletal dysplasias | 3/58 | 0.026 | 0.183 | 5.0 | 18.2 | *ADAMTSL2;RUNX3;SMAD7* |
| Endochondral Ossification with Skeletal Dysplasias | 4/64 | 0.005 | 0.101 | 6.1 | 32.0 | *ADAMTS5;ADAMTS1;RUNX3;BMP6* |
| Endochondral Ossification | 4/64 | 0.005 | 0.101 | 6.1 | 32.0 | *ADAMTS5;ADAMTS1;RUNX3;BMP6* |
| Primary focal segmental glomerulosclerosis | 3/72 | 0.045 | 0.275 | 4.0 | 12.3 | *LAMA5;AGRN;CLDN1* |
| Arrhythmogenic Cardiomyopathy | 5/74 | 0.001 | 0.058 | 6.7 | 44.3 | *CACNB1;TCF7L1;LAMA2;ACTN1;DMD* |
| Glioblastoma signaling pathways | 4/82 | 0.013 | 0.158 | 4.7 | 20.6 | *PDGFRA;CDK6;PRKCQ;FGFR2* |
| Small cell lung cancer | 4/96 | 0.021 | 0.183 | 4.0 | 15.3 | *LAMA5;CDK6;LAMA2;LAMA3* |
| Pathways Regulating Hippo Signaling | 4/98 | 0.023 | 0.183 | 3.9 | 14.7 | *PDGFRA;TCF7L1;PRKCQ;FGFR2* |
| Osteoblast Signaling | 3/14 | 0.000 | 0.038 | 25.0 | 193.4 | *PDGFRA;TNFRSF11B;FGF23* |
| Oncogenic Pathways as Metastatic Traits | 2/17 | 0.015 | 0.158 | 12.1 | 51.4 | *TCF7L1;TNC* |
| MFAP5 effects | 2/18 | 0.016 | 0.160 | 11.4 | 46.9 | *ACTN1;PRKCQ* |
| BMP Signaling in Eyelid Development | 3/20 | 0.001 | 0.058 | 16.1 | 107.5 | *FOXC1;PITX2;FGFR2* |
| FGF23 signaling | 2/22 | 0.024 | 0.183 | 9.1 | 34.0 | *FGF23;FGFR2* |
| Hfe effects | 2/7 | 0.002 | 0.086 | 36.5 | 219.6 | *BMP6;SMAD7* |
| Angiogenesis | 2/24 | 0.028 | 0.191 | 8.3 | 29.6 | *PDGFRA;FGFR2* |
| Ectoderm Differentiation | 10/138 | 0.000 | 0.001 | 7.3 | 93.6 | *TCF7L1;HDAC10;FZD4;BOC;HESX1;PAX3;TNFRSF11B;DMD;KIFC3;FGFR2* |
| Neural Crest Differentiation | 4/101 | 0.025 | 0.183 | 3.8 | 13.9 | *TCF7L1;HDAC10;PAX3;FGFR2* |
| Wnt signaling pathway and pluripotency | 4/102 | 0.026 | 0.183 | 3.7 | 13.7 | *TCF7L1;FZD4;LRRK2;PRKCQ* |
| ESC Pluripotency Pathways | 5/116 | 0.009 | 0.133 | 4.1 | 19.5 | *PDGFRA;FZD4;FGF23;FGFR2;SMAD7* |
| 22q11.2 copy number variation syndrome | 5/131 | 0.015 | 0.158 | 3.6 | 15.4 | *FOXC1;PAX3;PITX2;CLDN1;FGFR2* |
| TGF-beta Signaling Pathway | 5/132 | 0.015 | 0.158 | 3.6 | 15.1 | *CCNB2;SMURF2;TNC;SPTBN1;SMAD7* |
| Mesodermal commitment pathway | 5/147 | 0.023 | 0.183 | 3.2 | 12.2 | *FOXC1;TCF7L1;MEIS1;FZD4;PITX2* |
| Focal Adhesion | 7/198 | 0.006 | 0.101 | 3.4 | 17.2 | *PDGFRA;LAMA5;LAMA2;ACTN1;LAMA3;TNC;ARHGAP5* |
| MAPK Signaling Pathway | 8/246 | 0.006 | 0.101 | 3.1 | 16.0 | *DUSP4;CACNB1;PTPRR;LRRK2;RAPGEF2;FGF23;DUSP6;FGFR2* |
| PI3K-Akt signaling pathway | 8/340 | 0.034 | 0.224 | 2.2 | 7.5 | *PDGFRA;LAMA5;CDK6;LAMA2;LAMA3;TNC;FGF23;FGFR2* |

Note. The full enrichment analysis for genes shared between fish early domesticates and human groups with schizophrenia can be accessed here: <https://maayanlab.cloud/Enrichr/enrich?dataset=c68dc00d311e361026f7339bd11b73e8>

**Table S5**. Enrichment of Wikipathways associated with genes shared between early fish domesticates and human groups with Williams syndrome

| **Wikipathways** | **Overlap** | **P-value** | **Adjusted P-value** | **Odds Ratio** | **Combined Score** | **Genes** |
| --- | --- | --- | --- | --- | --- | --- |
| Ectoderm Differentiation | 9/138 | 1.95E-05 | 0.003 | 6.62 | 71.86 | *TCF7L1;HDAC10;FZD4;BOC;HESX1;PAX3;DMD;KIFC3;FGFR2* |
| Neural Crest Differentiation | 7/101 | 1.13E-04 | 0.009 | 7.02 | 63.74 | *TCF7L1;HDAC10;CDH2;PAX3;HES1;AXIN2;FGFR2* |
| Laminopathies on Wnt signaling | 4/35 | 5.33E-04 | 0.029 | 12.02 | 90.62 | *TCF7L1;ICMT;CDK6;HES1* |
| Mesodermal commitment pathway | 7/147 | 0.001 | 0.034 | 4.70 | 31.99 | *FOXC1;TCF7L1;MEIS1;FZD4;AXIN2;PITX2;TOX* |
| Arrhythmogenic Cardiomyopathy | 5/74 | 0.001 | 0.034 | 6.77 | 45.34 | *TCF7L1;CDH2;LAMA2;ACTN1;DMD* |
| BMP Signaling in Eyelid Development | 3/20 | 0.001 | 0.034 | 16.38 | 109.65 | *FOXC1;PITX2;FGFR2* |
| White fat cell differentiation | 3/32 | 0.005 | 0.079 | 9.59 | 51.04 | *TCF7L1;MECOM;ZNF423* |
| Hair Follicle Development | 3/32 | 0.005 | 0.079 | 9.59 | 51.04 | *LAMA5;PDGFRA;FGFR2* |

Note. The full enrichment analysis for genes shared between fish early domesticates and human groups with Williams syndrome can be accessed here: <https://maayanlab.cloud/Enrichr/enrich?dataset=b83024a0aa8fe2ff6492cb468c1d4ec0>

**Table S6**. Enrichment of Wikipathways associated with genes shared between early fish domesticates and human groups with autism spectrum disorders

| **Wikipathways** | **Overlap** | **P-value** | **Adjusted P-value** | **Odds Ratio** | **Combined Score** | **Genes** |
| --- | --- | --- | --- | --- | --- | --- |
| Ectoderm Differentiation | 7/138 | 1.15E-04 | 0.012 | 6.98 | 63.29 | *TFAP2A;BOC;HESX1;DMD;KIFC3;FGFR2;GLI3* |
| Alpha 6 Beta 4 signaling pathway | 4/33 | 1.28E-04 | 0.012 | 17.75 | 159.02 | *LAMA5;LAMA2;LAMA3;PIK3R1* |
| Metastatic brain tumor | 2/6 | 9.11E-04 | 0.046 | 63.58 | 445.14 | *CDK6;PIK3R1* |
| Small cell lung cancer | 5/96 | 9.90E-04 | 0.046 | 7.09 | 49.07 | *LAMA5;CDK6;LAMA2;LAMA3;PIK3R1* |
| Neural Crest Differentiation | 5/101 | 0.001 | 0.046 | 6.72 | 44.97 | *TFAP2A;TFAP2B;CDH2;AXIN2;FGFR2* |
| Hair Follicle Development | 3/32 | 0.002 | 0.063 | 13.22 | 81.99 | *LAMA5;PDGFRA;FGFR2* |
| Arrhythmogenic Cardiomyopathy | 4/74 | 0.003 | 0.074 | 7.34 | 43.10 | *CDH2;LAMA2;ACTN1;DMD* |
| Glioblastoma signaling pathways | 4/82 | 0.004 | 0.093 | 6.58 | 36.23 | *PDGFRA;CDK6;PIK3R1;FGFR2* |
| Focal Adhesion | 6/198 | 0.005 | 0.093 | 4.04 | 21.45 | *PDGFRA;LAMA5;LAMA2;ACTN1;LAMA3;PIK3R1* |
| Hedgehog Signaling Pathway | 3/44 | 0.005 | 0.093 | 9.35 | 49.43 | *SMURF2;BOC;GLI3* |
| Pathways Regulating Hippo Signaling | 4/98 | 0.008 | 0.128 | 5.46 | 26.61 | *PDGFRA;CDH2;PRKAR1B;FGFR2* |
| Endometrial cancer | 3/63 | 0.014 | 0.166 | 6.38 | 27.43 | *AXIN2;PIK3R1;FGFR2* |
| GDNF/RET signaling axis | 2/23 | 0.014 | 0.166 | 12.10 | 51.59 | *HSPB11;GLI3* |
| Endochondral Ossification with Skeletal Dysplasias | 3/64 | 0.014 | 0.166 | 6.28 | 26.71 | *RUNX3;GLI3;BMP6* |
| Endochondral Ossification | 3/64 | 0.014 | 0.166 | 6.28 | 26.71 | *RUNX3;GLI3;BMP6* |
| Hippo-Merlin Signaling Dysregulation | 4/120 | 0.015 | 0.166 | 4.42 | 18.50 | *PDGFRA;CDH2;PRKAR1B;FGFR2* |
| Angiogenesis | 2/24 | 0.015 | 0.166 | 11.55 | 48.30 | *PDGFRA;FGFR2* |
| PI3K-Akt signaling pathway | 7/340 | 0.019 | 0.186 | 2.72 | 10.82 | *PDGFRA;LAMA5;CDK6;LAMA2;LAMA3;PIK3R1;FGFR2* |
| Primary focal segmental glomerulosclerosis | 3/72 | 0.019 | 0.186 | 5.55 | 21.87 | *LAMA5;CDH2;AGRN* |
| Head and Neck Squamous Cell Carcinoma | 3/73 | 0.020 | 0.186 | 5.47 | 21.36 | *CDK6;PIK3R1;FGFR2* |
| Tumor suppressor activity of SMARCB1 | 2/31 | 0.025 | 0.219 | 8.76 | 32.37 | *CDK6;GLI3* |
| EGFR Tyrosine Kinase Inhibitor Resistance | 3/84 | 0.029 | 0.241 | 4.72 | 16.73 | *PDGFRA;PIK3R1;FGFR2* |
| Acute viral myocarditis | 3/86 | 0.031 | 0.241 | 4.61 | 16.04 | *LAMA2;DMD;PIK3R1* |
| Regulation of Actin Cytoskeleton | 4/150 | 0.031 | 0.241 | 3.50 | 12.14 | *PDGFRA;ACTN1;PIK3R1;FGFR2* |
| Focal Adhesion-PI3K-Akt-mTOR-signaling | 6/303 | 0.034 | 0.242 | 2.60 | 8.82 | *PDGFRA;LAMA5;LAMA2;LAMA3;PIK3R1;FGFR2* |
| Breast cancer pathway | 4/154 | 0.034 | 0.242 | 3.41 | 11.53 | *CDK6;PIK3R1;AXIN2;ESR1* |
| Regulation of GABA neurotransmission | 2/38 | 0.036 | 0.248 | 7.05 | 23.41 | *PIK3R1;GABRD* |
| LncRNA involvement in Wnt signaling | 3/94 | 0.038 | 0.254 | 4.20 | 13.68 | *TFAP2A;CDK6;AXIN2* |
| Splicing factor NOVA regulated synaptic proteins | 2/42 | 0.043 | 0.277 | 6.35 | 19.90 | *CDH2;AGRN* |
| Genes controlling nephrogenesis | 2/43 | 0.045 | 0.280 | 6.19 | 19.15 | *GLI3;FGFR2* |

Note. The full enrichment analysis for genes shared between fish early domesticates and human groups with autism spectrum disorders can be accessed here: <https://maayanlab.cloud/Enrichr/enrich?dataset=c5b8b850d070c8fa90b809675170affd>

**Table S7**. Enrichment of Gene Ontology (GO) Biological Process (BP) terms associated with genes shared between early fish domesticates and human groups with schizophrenia

| **Biological process GO** | **ID** | **Overlap** | **P-value** | **Adjusted P-value** | **Odds Ratio** | **Combined Score** | **Genes** |
| --- | --- | --- | --- | --- | --- | --- | --- |
| extracellular structure organization | GO:0043062 | 12/216 | 4.89E-06 | 0.003 | 5.6 | 68.0 | *ADAMTS5;LAMA5;LAMA2;ADAMTS1;ADAMTS18;ADAMTS17;ADAMTSL2;LAMA3;TNC;AGRN;ADAMTS9;ADAMTS6* |
| external encapsulating structure organization | GO:0045229 | 12/217 | 5.13E-06 | 0.003 | 5.5 | 67.4 | *ADAMTS5;LAMA5;LAMA2;ADAMTS1;ADAMTS18;ADAMTS17;ADAMTSL2;LAMA3;TNC;AGRN;ADAMTS9;ADAMTS6* |
| extracellular matrix organization | GO:0030198 | 12/300 | 1.23E-04 | 0.034 | 3.9 | 35.3 | *ADAMTS5;LAMA5;LAMA2;ADAMTS1;ADAMTS18;ADAMTS17;ADAMTSL2;LAMA3;TNC;AGRN;ADAMTS9;ADAMTS6* |
| neuromuscular junction development | GO:0007528 | 4/24 | 1.25E-04 | 0.034 | 18.4 | 165.2 | *CACNB1;MUSK;LRRK2;AGRN* |
| MAPK cascade | GO:0000165 | 12/303 | 1.35E-04 | 0.034 | 3.9 | 34.6 | *PSMB7;PDGFRA;PSMB4;DLG3;LRRK2;RAPGEF2;PRKCQ;FGF23;SPTBN1;DUSP6;FGFR2;PTPN3* |
| morphogenesis of an epithelium | GO:0002009 | 4/30 | 3.07E-04 | 0.064 | 14.1 | 114.4 | *LAMA5;LAMA3;TBX18;FGFR2* |
| regulation of membrane depolarization | GO:0003254 | 3/18 | 9.36E-04 | 0.160 | 18.3 | 127.6 | *LRRK2;SMAD7;PTPN3* |
| regulation of plasma membrane bounded cell projection assembly | GO:0120032 | 5/70 | 0.001 | 0.160 | 7.1 | 48.8 | *TBC1D2;ATP8B1;TBC1D8;TBC1D22A;TBC1D22B* |
| regulation of morphogenesis of a branching structure | GO:0060688 | 2/5 | 0.001 | 0.162 | 60.8 | 410.3 | *LRRK2;FGFR2* |
| negative regulation of ERK1 and ERK2 cascade | GO:0070373 | 4/50 | 0.002 | 0.212 | 8.0 | 48.9 | *DUSP4;PTPRR;DUSP26;DUSP6* |
| positive regulation of phospholipase activity | GO:0010518 | 3/24 | 0.002 | 0.212 | 13.1 | 79.9 | *PDGFRA;ESR1;FGFR2* |
| glandular epithelial cell development | GO:0002068 | 2/7 | 0.002 | 0.212 | 36.5 | 219.6 | *CDK6;BMP6* |
| type B pancreatic cell development | GO:0003323 | 2/7 | 0.002 | 0.212 | 36.5 | 219.6 | *CDK6;BMP6* |
| ventricular cardiac muscle tissue development | GO:0003229 | 3/25 | 0.002 | 0.212 | 12.5 | 74.8 | *ADAMTS9;FGFR2;SMAD7* |
| positive regulation of GTPase activity | GO:0043547 | 8/214 | 0.003 | 0.212 | 3.6 | 21.5 | *RABGAP1L;TBC1D2;TBC1D8;RAPGEF2;ARAP1;AGRN;TBC1D22A;TBC1D22B* |
| regulation of anion transport | GO:0044070 | 2/8 | 0.003 | 0.225 | 30.4 | 174.5 | *ATP8B1;FGF23* |
| forebrain neuron development | GO:0021884 | 2/8 | 0.003 | 0.225 | 30.4 | 174.5 | *RAPGEF2;FGFR2* |
| receptor clustering | GO:0043113 | 3/28 | 0.003 | 0.225 | 11.0 | 62.2 | *MUSK;DLG3;AGRN* |
| eye development | GO:0001654 | 4/58 | 0.004 | 0.225 | 6.8 | 38.0 | *FOXC1;MEIS1;ADAMTS18;PITX2* |
| neuronal ion channel clustering | GO:0045161 | 2/9 | 0.004 | 0.225 | 26.0 | 143.2 | *KCNIP2;AGRN* |
| embryonic digestive tract morphogenesis | GO:0048557 | 2/9 | 0.004 | 0.225 | 26.0 | 143.2 | *PDGFRA;FGFR2* |
| type B pancreatic cell differentiation | GO:0003309 | 2/9 | 0.004 | 0.225 | 26.0 | 143.2 | *CDK6;BMP6* |
| maintenance of blood-brain barrier | GO:0035633 | 3/30 | 0.004 | 0.225 | 10.2 | 55.5 | *LAMA2;DMD;CLDN1* |
| protein dephosphorylation | GO:0006470 | 6/139 | 0.004 | 0.225 | 4.2 | 22.7 | *DUSP4;PTPRR;EYA2;DUSP26;DUSP6;PTPN3* |
| embryonic skeletal system morphogenesis | GO:0048704 | 3/31 | 0.005 | 0.232 | 9.8 | 52.6 | *PDGFRA;FGFR2;HOXA4* |
| regulation of T-helper 17 type immune response | GO:2000316 | 2/10 | 0.005 | 0.244 | 22.8 | 120.4 | *PRKCQ;SMAD7* |
| regulation of cilium assembly | GO:1902017 | 4/64 | 0.005 | 0.247 | 6.1 | 32.0 | *TBC1D2;TBC1D8;TBC1D22A;TBC1D22B* |
| activation of GTPase activity | GO:0090630 | 5/105 | 0.006 | 0.256 | 4.6 | 23.5 | *RABGAP1L;TBC1D2;TBC1D8;TBC1D22A;TBC1D22B* |
| cell-substrate junction assembly | GO:0007044 | 3/34 | 0.006 | 0.256 | 8.8 | 45.2 | *ACTN1;LAMA3;SORBS1* |
| positive regulation of alcohol biosynthetic process | GO:1902932 | 2/11 | 0.006 | 0.256 | 20.2 | 103.1 | *SCAP;BMP6* |
| morphogenesis of a polarized epithelium | GO:0001738 | 2/12 | 0.007 | 0.295 | 18.2 | 89.6 | *LAMA5;LAMA3* |
| regulation of Wnt signaling pathway | GO:0030111 | 5/111 | 0.008 | 0.295 | 4.3 | 21.2 | *TCF7L1;LRRK2;ESR1;CXXC4;FGFR2* |
| negative regulation of epithelial cell proliferation | GO:0050680 | 4/72 | 0.008 | 0.305 | 5.4 | 26.0 | *CDK6;RUNX3;XDH;FGFR2* |
| positive regulation of kinase activity | GO:0033674 | 5/114 | 0.008 | 0.310 | 4.2 | 20.1 | *PDGFRA;MUSK;AGAP2;RAPGEF2;FGFR2* |
| odontogenesis | GO:0042476 | 3/39 | 0.009 | 0.317 | 7.6 | 36.0 | *FOXC1;PITX2;FGFR2* |
| regulation of sodium ion transmembrane transporter activity | GO:2000649 | 3/40 | 0.009 | 0.326 | 7.4 | 34.5 | *NETO1;DMD;PTPN3* |
| regulation of vasculogenesis | GO:2001212 | 2/14 | 0.010 | 0.326 | 15.2 | 70.0 | *RAPGEF2;XDH* |
| regulation of organelle assembly | GO:1902115 | 4/77 | 0.010 | 0.326 | 5.0 | 23.1 | *TBC1D2;TBC1D8;TBC1D22A;TBC1D22B* |
| visual system development | GO:0150063 | 3/41 | 0.010 | 0.326 | 7.2 | 33.1 | *FOXC1;MEIS1;ADAMTS18* |
| skeletal system morphogenesis | GO:0048705 | 3/42 | 0.011 | 0.337 | 7.0 | 31.8 | *PDGFRA;FGFR2;HOXA4* |
| regulation of phospholipase C activity | GO:1900274 | 2/15 | 0.011 | 0.337 | 14.0 | 62.7 | *PDGFRA;ESR1* |
| embryonic cranial skeleton morphogenesis | GO:0048701 | 2/15 | 0.011 | 0.337 | 14.0 | 62.7 | *PDGFRA;FGFR2* |
| negative regulation of cell cycle | GO:0045786 | 4/80 | 0.012 | 0.337 | 4.8 | 21.5 | *FOXC1;CDK6;RUNX3;PTPN3* |
| embryonic organ morphogenesis | GO:0048562 | 3/44 | 0.012 | 0.342 | 6.7 | 29.4 | *PDGFRA;FGFR2;HOXA4* |
| negative regulation of platelet activation | GO:0010544 | 2/16 | 0.013 | 0.342 | 13.0 | 56.6 | *PDGFRA;ADAMTS18* |
| regulation of monocyte differentiation | GO:0045655 | 2/16 | 0.013 | 0.342 | 13.0 | 56.6 | *CDK6;HOXA7* |
| digestive tract morphogenesis | GO:0048546 | 2/16 | 0.013 | 0.342 | 13.0 | 56.6 | *PDGFRA;FGFR2* |
| regulation of osteoblast differentiation | GO:0045667 | 4/83 | 0.013 | 0.342 | 4.6 | 20.1 | *CDK6;FGF23;BMP6;FGFR2* |
| Wnt signaling pathway, planar cell polarity pathway | GO:0060071 | 4/85 | 0.014 | 0.350 | 4.5 | 19.2 | *PSMB7;PSMB4;SMURF2;FZD4* |
| positive regulation of canonical Wnt signaling pathway | GO:0090263 | 5/130 | 0.014 | 0.350 | 3.7 | 15.6 | *PSMB7;PSMB4;SMURF2;LRRK2;FGFR2* |
| mesonephric tubule development | GO:0072164 | 2/17 | 0.015 | 0.350 | 12.1 | 51.4 | *FOXC1;FGFR2* |
| vasculature development | GO:0001944 | 2/17 | 0.015 | 0.350 | 12.1 | 51.4 | *PDGFRA;SOX18* |
| regulation of establishment of planar polarity | GO:0090175 | 4/88 | 0.016 | 0.363 | 4.4 | 18.1 | *PSMB7;PSMB4;SMURF2;FZD4* |
| regulation of ERK1 and ERK2 cascade | GO:0070372 | 7/238 | 0.016 | 0.363 | 2.8 | 11.5 | *DUSP4;PDGFRA;PTPRR;RAPGEF2;DUSP26;DUSP6;FGFR2* |
| regulation of platelet aggregation | GO:0090330 | 2/18 | 0.016 | 0.363 | 11.4 | 46.9 | *ADAMTS18;PRKCQ* |
| embryonic heart tube development | GO:0035050 | 2/18 | 0.016 | 0.363 | 11.4 | 46.9 | *SOX18;PITX2* |
| regulation of cation channel activity | GO:2001257 | 4/89 | 0.017 | 0.363 | 4.3 | 17.7 | *CACNB1;NETO1;DLG3;DMD* |
| axonogenesis | GO:0007409 | 7/240 | 0.017 | 0.363 | 2.8 | 11.3 | *EFNB1;LAMA5;LAMA2;LAMA3;PRKCQ;SPTBN1;FGFR2* |
| protein complex oligomerization | GO:0051259 | 4/90 | 0.017 | 0.363 | 4.3 | 17.3 | *EHD4;C9;ADCY8;CLDN1* |
| regulation of cellular component movement | GO:0051270 | 3/50 | 0.017 | 0.363 | 5.8 | 23.6 | *CDK6;ACTN1;ARAP1* |
| ureteric bud development | GO:0001657 | 2/19 | 0.018 | 0.367 | 10.7 | 43.0 | *FOXC1;FGFR2* |
| mesenchymal cell differentiation | GO:0048762 | 3/51 | 0.018 | 0.367 | 5.7 | 22.8 | *FOXC1;FBXL17;FGFR2* |
| Wnt signaling pathway | GO:0016055 | 4/92 | 0.018 | 0.367 | 4.2 | 16.6 | *TCF7L1;FZD4;LRRK2;STRN* |
| regulation of epithelial cell proliferation | GO:0050678 | 4/93 | 0.019 | 0.374 | 4.1 | 16.3 | *CDK6;RUNX3;BMP6;FGFR2* |
| negative regulation of MAPK cascade | GO:0043409 | 4/94 | 0.020 | 0.378 | 4.1 | 16.0 | *DUSP4;PTPRR;DUSP26;DUSP6* |
| embryonic digestive tract development | GO:0048566 | 2/20 | 0.020 | 0.378 | 10.1 | 39.6 | *PDGFRA;FGFR2* |
| negative regulation of peptidyl-serine phosphorylation | GO:0033137 | 2/21 | 0.022 | 0.382 | 9.6 | 36.6 | *DMD;SMAD7* |
| focal adhesion assembly | GO:0048041 | 2/21 | 0.022 | 0.382 | 9.6 | 36.6 | *ACTN1;SORBS1* |
| sensory organ development | GO:0007423 | 3/56 | 0.023 | 0.382 | 5.2 | 19.4 | *FOXC1;MEIS1;ADAMTS18* |
| negative regulation of myeloid cell differentiation | GO:0045638 | 2/22 | 0.024 | 0.382 | 9.1 | 34.0 | *MEIS1;CDK6* |
| peptidyl-threonine dephosphorylation | GO:0035970 | 2/22 | 0.024 | 0.382 | 9.1 | 34.0 | *DUSP4;DUSP6* |
| regulation of platelet activation | GO:0010543 | 2/22 | 0.024 | 0.382 | 9.1 | 34.0 | *PDGFRA;PRKCQ* |
| regulation of voltage-gated calcium channel activity | GO:1901385 | 2/22 | 0.024 | 0.382 | 9.1 | 34.0 | *CACNB1;DMD* |
| proteasomal ubiquitin-independent protein catabolic process | GO:0010499 | 2/22 | 0.024 | 0.382 | 9.1 | 34.0 | *PSMB7;PSMB4* |
| axon guidance | GO:0007411 | 6/203 | 0.025 | 0.382 | 2.8 | 10.4 | *EFNB1;LAMA5;LAMA2;LAMA3;PRKCQ;SPTBN1* |
| regulation of pathway-restricted SMAD protein phosphorylation | GO:0060393 | 3/58 | 0.026 | 0.382 | 5.0 | 18.2 | *BMP6;BMP15;SMAD7* |
| regulation of protein kinase B signaling | GO:0051896 | 6/207 | 0.027 | 0.382 | 2.7 | 9.9 | *PDGFRA;STRN;XDH;FGF23;ESR1;FGFR2* |
| dephosphorylation | GO:0016311 | 5/153 | 0.027 | 0.382 | 3.1 | 11.2 | *DUSP4;PTPRR;DUSP26;DUSP6;PTPN3* |
| positive regulation of Wnt signaling pathway | GO:0030177 | 5/153 | 0.027 | 0.382 | 3.1 | 11.2 | *PSMB7;PSMB4;SMURF2;LRRK2;FGFR2* |
| response to organic cyclic compound | GO:0014070 | 3/60 | 0.028 | 0.382 | 4.8 | 17.2 | *LRRK2;ESR1;STRN3* |
| regulation of cellular response to transforming growth factor beta stimulus | GO:1903844 | 2/24 | 0.028 | 0.382 | 8.3 | 29.6 | *SMURF2;SMAD7* |
| inner ear morphogenesis | GO:0042472 | 2/24 | 0.028 | 0.382 | 8.3 | 29.6 | *TBX18;FGFR2* |
| lipid translocation | GO:0034204 | 2/24 | 0.028 | 0.382 | 8.3 | 29.6 | *ATP8B1;ATP11A* |
| regulation of cell differentiation | GO:0045595 | 5/156 | 0.029 | 0.382 | 3.0 | 10.8 | *ZFHX3;CDK6;RUNX3;FGFR2;SMAD7* |
| in utero embryonic development | GO:0001701 | 2/25 | 0.030 | 0.382 | 7.9 | 27.7 | *PTPRR;FGFR2* |
| negative regulation of transcription, DNA-templated | GO:0045892 | 17/948 | 0.032 | 0.382 | 1.7 | 5.9 | *FOXC1;ZFHX3;L3MBTL2;HDAC10;SMURF2;ATP8B1;DUSP26;RUNX3;ESR1;BMP6;SMAD7;SOX18;PAX9;HESX1;HOXA7;FGFR2;SAMD11* |
| positive regulation of protein kinase B signaling | GO:0051897 | 5/161 | 0.032 | 0.382 | 2.9 | 10.1 | *PDGFRA;STRN;FGF23;ESR1;FGFR2* |
| endothelial cell development | GO:0001885 | 2/26 | 0.033 | 0.382 | 7.6 | 26.0 | *SOX18;RAPGEF2* |
| positive regulation of filopodium assembly | GO:0051491 | 2/26 | 0.033 | 0.382 | 7.6 | 26.0 | *ARAP1;AGRN* |
| transmembrane receptor protein tyrosine kinase signaling pathway | GO:0007169 | 9/404 | 0.034 | 0.382 | 2.1 | 7.1 | *PDGFRA;EFNB1;FOXC1;TIA1;MUSK;RAPGEF2;SORBS1;FGF23;FGFR2* |
| BMP signaling pathway | GO:0030509 | 3/65 | 0.034 | 0.382 | 4.4 | 14.9 | *BMP6;BMP15;SMAD7* |
| positive regulation of fibroblast proliferation | GO:0048146 | 2/28 | 0.037 | 0.382 | 7.0 | 23.0 | *PDGFRA;CDK6* |
| positive regulation of phosphatidylinositol 3-kinase activity | GO:0043552 | 2/28 | 0.037 | 0.382 | 7.0 | 23.0 | *PDGFRA;AGAP2* |
| canonical Wnt signaling pathway | GO:0060070 | 3/68 | 0.039 | 0.382 | 4.2 | 13.7 | *TCF7L1;FZD4;LRRK2* |
| negative regulation of transforming growth factor beta receptor signaling pathway | GO:0030512 | 3/68 | 0.039 | 0.382 | 4.2 | 13.7 | *SMURF2;ADAMTSL2;SMAD7* |
| regulation of myeloid cell differentiation | GO:0045637 | 3/68 | 0.039 | 0.382 | 4.2 | 13.7 | *MEIS1;CDK6;PRKCQ* |
| negative regulation of transcription by RNA polymerase II | GO:0000122 | 13/684 | 0.039 | 0.382 | 1.8 | 5.8 | *FOXC1;ZFHX3;HDAC10;SMURF2;DUSP26;RUNX3;ESR1;BMP6;SMAD7;SOX18;HESX1;HOXA7;FGFR2* |
| regulation of calcium ion transmembrane transporter activity | GO:1901019 | 2/29 | 0.040 | 0.382 | 6.7 | 21.7 | *CACNB1;DMD* |
| negative regulation of myeloid leukocyte differentiation | GO:0002762 | 2/29 | 0.040 | 0.382 | 6.7 | 21.7 | *CDK6;HOXA7* |
| establishment of endothelial barrier | GO:0061028 | 2/29 | 0.040 | 0.382 | 6.7 | 21.7 | *SOX18;RAPGEF2* |
| artery morphogenesis | GO:0048844 | 2/30 | 0.042 | 0.382 | 6.5 | 20.5 | *ADAMTS9;SMAD7* |
| cardiac muscle cell action potential | GO:0086001 | 2/30 | 0.042 | 0.382 | 6.5 | 20.5 | *KCNIP2;DMD* |
| regulation of striated muscle contraction | GO:0006942 | 2/30 | 0.042 | 0.382 | 6.5 | 20.5 | *DMD;SMAD7* |
| cellular response to BMP stimulus | GO:0071773 | 3/71 | 0.043 | 0.382 | 4.0 | 12.7 | *BMP6;BMP15;SMAD7* |
| regulation of hematopoietic stem cell differentiation | GO:1902036 | 3/71 | 0.043 | 0.382 | 4.0 | 12.7 | *PSMB7;FOXC1;PSMB4* |
| cell morphogenesis involved in differentiation | GO:0000904 | 3/72 | 0.045 | 0.382 | 4.0 | 12.3 | *ATRN;LAMA5;ACTN1* |
| negative regulation of osteoblast differentiation | GO:0045668 | 2/31 | 0.045 | 0.382 | 6.3 | 19.5 | *CDK6;FGF23* |
| phospholipid translocation | GO:0045332 | 2/31 | 0.045 | 0.382 | 6.3 | 19.5 | *ATP8B1;ATP11A* |
| regulation of transmembrane receptor protein serine/threonine kinase signaling pathway | GO:0090092 | 2/31 | 0.045 | 0.382 | 6.3 | 19.5 | *SMURF2;SMAD7* |
| non-canonical Wnt signaling pathway | GO:0035567 | 4/124 | 0.048 | 0.382 | 3.0 | 9.3 | *PSMB7;PSMB4;SMURF2;FZD4* |
| cardiac muscle tissue morphogenesis | GO:0055008 | 2/32 | 0.048 | 0.382 | 6.1 | 18.5 | *FGFR2;SMAD7* |
| regulation of intracellular estrogen receptor signaling pathway | GO:0033146 | 2/32 | 0.048 | 0.382 | 6.1 | 18.5 | *ESR1;STRN3* |
| metal ion export | GO:0070839 | 2/32 | 0.048 | 0.382 | 6.1 | 18.5 | *ATP7B;KCNIP2* |
| regulation of hematopoietic progenitor cell differentiation | GO:1901532 | 3/75 | 0.049 | 0.382 | 3.8 | 11.4 | *PSMB7;FOXC1;PSMB4* |
| fibroblast growth factor receptor signaling pathway | GO:0008543 | 3/75 | 0.049 | 0.382 | 3.8 | 11.4 | *TIA1;FGF23;FGFR2* |
| regulation of transcription from RNA polymerase II promoter in response to hypoxia | GO:0061418 | 3/75 | 0.049 | 0.382 | 3.8 | 11.4 | *PSMB7;PSMB4;EGLN3* |
| integrin-mediated signaling pathway | GO:0007229 | 3/75 | 0.049 | 0.382 | 3.8 | 11.4 | *LAMA5;ADAMTS1;LAMA3* |
| synapse organization | GO:0050808 | 4/126 | 0.050 | 0.382 | 3.0 | 9.0 | *CACNB1;MUSK;LRRK2;AGRN* |

Note. The full enrichment analysis for genes shared between fish early domesticates and human groups with schizophrenia can be accessed here: <https://maayanlab.cloud/Enrichr/enrich?dataset=c68dc00d311e361026f7339bd11b73e8>

**Table S8**. Enrichment of Gene Ontology (GO) Biological Process (BP) terms associated with genes shared between early fish domesticates and human groups with Williams syndrome

| **Biological process GO** | **ID** | **Overlap** | **P-value** | **Adjusted P-value** | **Odds Ratio** | **Combined Score** | **Genes** |
| --- | --- | --- | --- | --- | --- | --- | --- |
| extracellular structure organization | GO:0043062 | 12/216 | 4.25E-06 | 0.003 | 5.6 | 69.8 | *ADAMTS5;LAMA5;LAMA2;ADAMTS1;ADAMTS18;ADAMTS17;ADAMTSL2;LAMA3;TNC;AGRN;ADAMTS9;ADAMTS6* |
| external encapsulating structure organization | GO:0045229 | 12/217 | 4.45E-06 | 0.003 | 5.6 | 69.2 | *ADAMTS5;LAMA5;LAMA2;ADAMTS1;ADAMTS18;ADAMTS17;ADAMTSL2;LAMA3;TNC;AGRN;ADAMTS9;ADAMTS6* |
| extracellular matrix organization | GO:0030198 | 13/300 | 2.46E-05 | 0.011 | 4.4 | 46.2 | *LAMA5;LAMA2;LAMA3;TNC;ADAMTS5;P4HA2;ADAMTS1;ADAMTS18;ADAMTS17;ADAMTSL2;AGRN;ADAMTS9;ADAMTS6* |
| glandular epithelial cell development | GO:0002068 | 3/7 | 4.21E-05 | 0.011 | 69.6 | 701.7 | *CDK6;CDH2;BMP6* |
| type B pancreatic cell development | GO:0003323 | 3/7 | 4.21E-05 | 0.011 | 69.6 | 701.7 | *CDK6;CDH2;BMP6* |
| type B pancreatic cell differentiation | GO:0003309 | 3/9 | 9.95E-05 | 0.022 | 46.4 | 427.9 | *CDK6;CDH2;BMP6* |
| neuromuscular junction development | GO:0007528 | 4/24 | 1.19E-04 | 0.022 | 18.6 | 168.5 | *MUSK;LRRK2;LRP4;AGRN* |
| receptor clustering | GO:0043113 | 4/28 | 2.21E-04 | 0.036 | 15.5 | 130.8 | *MUSK;DLG3;LRP4;AGRN* |
| morphogenesis of an epithelium | GO:0002009 | 4/30 | 2.91E-04 | 0.043 | 14.3 | 116.7 | *LAMA5;LAMA3;TBX18;FGFR2* |
| negative regulation of multicellular organismal process | GO:0051241 | 9/214 | 5.49E-04 | 0.071 | 4.2 | 31.2 | *ADAMTS5;PTPRR;NFIB;LRP4;HOXA7;IL17D;ZNF423;N4BP1;TBX18* |
| anterior/posterior pattern specification | GO:0009952 | 5/63 | 5.93E-04 | 0.071 | 8.1 | 59.9 | *HES6;BHLHE41;HES1;HOXA7;HOXA4* |
| gliogenesis | GO:0042063 | 3/17 | 7.55E-04 | 0.082 | 19.9 | 143.0 | *CDK6;CDH2;NFIB* |
| odontogenesis | GO:0042476 | 4/39 | 8.09E-04 | 0.082 | 10.6 | 75.8 | *FOXC1;AXIN2;PITX2;FGFR2* |
| regulation of morphogenesis of a branching structure | GO:0060688 | 2/5 | 0.001 | 0.107 | 61.6 | 417.8 | *LRRK2;FGFR2* |
| embryonic organ morphogenesis | GO:0048562 | 4/44 | 0.001 | 0.113 | 9.3 | 62.0 | *PDGFRA;HES1;FGFR2;HOXA4* |
| positive regulation of kinase activity | GO:0033674 | 6/114 | 0.001 | 0.124 | 5.2 | 33.8 | *PDGFRA;MUSK;AGAP2;SPDYA;RAPGEF2;FGFR2* |
| MAPK cascade | GO:0000165 | 10/303 | 0.002 | 0.136 | 3.2 | 20.5 | *PDGFRA;ICMT;DLG3;LRRK2;RAPGEF2;FGF23;SPTBN1;DUSP6;FGFR2;PTPN3* |
| negative regulation of ERK1 and ERK2 cascade | GO:0070373 | 4/50 | 0.002 | 0.146 | 8.1 | 50.1 | *DUSP4;PTPRR;DUSP26;DUSP6* |
| endosome to lysosome transport via multivesicular body sorting pathway | GO:0032510 | 2/7 | 0.002 | 0.146 | 37.0 | 223.8 | *VPS4B;LYST* |
| positive regulation of synaptic plasticity | GO:0031915 | 2/7 | 0.002 | 0.146 | 37.0 | 223.8 | *ADCY8;CPLX2* |
| protein localization to cytoplasmic stress granule | GO:1903608 | 2/7 | 0.002 | 0.146 | 37.0 | 223.8 | *TIA1;DHX9* |
| positive regulation of protein phosphorylation | GO:0001934 | 11/371 | 0.002 | 0.146 | 2.9 | 17.4 | *MARCO;MUSK;LRRK2;AGAP2;SPDYA;RAPGEF2;HCLS1;AXIN2;FGF23;BMP6;BMP15* |
| positive regulation of binding | GO:0051099 | 5/90 | 0.003 | 0.158 | 5.5 | 32.0 | *FOXC1;DHX9;LRRK2;RAPGEF2;HES1* |
| regulation of anion transport | GO:0044070 | 2/8 | 0.003 | 0.158 | 30.8 | 177.8 | *ATP8B1;FGF23* |
| peptidyl-proline hydroxylation to 4-hydroxy-L-proline | GO:0018401 | 2/8 | 0.003 | 0.158 | 30.8 | 177.8 | *P4HA2;P4HTM* |
| forebrain neuron development | GO:0021884 | 2/8 | 0.003 | 0.158 | 30.8 | 177.8 | *RAPGEF2;FGFR2* |
| positive regulation of fibroblast proliferation | GO:0048146 | 3/28 | 0.003 | 0.162 | 11.1 | 63.5 | *PDGFRA;CDK6;DHX9* |
| hemopoiesis | GO:0030097 | 5/94 | 0.004 | 0.162 | 5.2 | 29.6 | *MEIS1;MECOM;TOX;RUNX3;FGFR2* |
| eye development | GO:0001654 | 4/58 | 0.004 | 0.162 | 6.9 | 38.9 | *FOXC1;MEIS1;ADAMTS18;PITX2* |
| embryonic digestive tract morphogenesis | GO:0048557 | 2/9 | 0.004 | 0.172 | 26.4 | 146.0 | *PDGFRA;FGFR2* |
| protein dephosphorylation | GO:0006470 | 6/139 | 0.004 | 0.172 | 4.2 | 23.3 | *DUSP4;PTPRR;SBF1;DUSP26;DUSP6;PTPN3* |
| regulation of ERK1 and ERK2 cascade | GO:0070372 | 8/238 | 0.004 | 0.179 | 3.3 | 17.7 | *DUSP4;MARCO;PDGFRA;PTPRR;RAPGEF2;DUSP26;DUSP6;FGFR2* |
| embryonic skeletal system morphogenesis | GO:0048704 | 3/31 | 0.004 | 0.179 | 9.9 | 53.8 | *PDGFRA;FGFR2;HOXA4* |
| negative regulation of nucleic acid-templated transcription | GO:1903507 | 12/464 | 0.005 | 0.184 | 2.5 | 13.5 | *L3MBTL2;HDAC10;MECOM;SMURF2;ATP8B1;PAX9;BHLHE41;HES1;HOXA7;ZNF423;NPAS1;SAMD11* |

Note. The full enrichment analysis for genes shared between fish early domesticates and human groups with Williams syndrome can be accessed here: <https://maayanlab.cloud/Enrichr/enrich?dataset=b83024a0aa8fe2ff6492cb468c1d4ec0>

**Table S9**. Enrichment of Gene Ontology (GO) Biological Process (BP) terms associated with genes shared between early fish domesticates and human groups with autism spectrum disorders

| **Biological process GO** | **ID** | **Overlap** | **P-value** | **Adjusted P-value** | **Odds Ratio** | **Combined Score** | **Genes** |
| --- | --- | --- | --- | --- | --- | --- | --- |
| renal system development | GO:0072001 | 6/57 | 5.75E-06 | 0.005 | 15.3 | 184.8 | *TFAP2A;TFAP2B;LRP4;SULF1;HSPB11;TBX18* |
| glandular epithelial cell development | GO:0002068 | 3/7 | 1.65E-05 | 0.005 | 96.0 | 1056.8 | *CDK6;CDH2;BMP6* |
| type B pancreatic cell development | GO:0003323 | 3/7 | 1.65E-05 | 0.005 | 96.0 | 1056.8 | *CDK6;CDH2;BMP6* |
| type B pancreatic cell differentiation | GO:0003309 | 3/9 | 3.92E-05 | 0.009 | 64.0 | 649.2 | *CDK6;CDH2;BMP6* |
| morphogenesis of an epithelium | GO:0002009 | 4/30 | 8.75E-05 | 0.017 | 19.8 | 185.0 | *LAMA5;LAMA3;TBX18;FGFR2* |
| embryonic cranial skeleton morphogenesis | GO:0048701 | 3/15 | 2.05E-04 | 0.029 | 32.0 | 271.6 | *TFAP2A;PDGFRA;FGFR2* |
| kidney development | GO:0001822 | 5/70 | 2.31E-04 | 0.029 | 9.9 | 83.3 | *TFAP2A;TFAP2B;LRP4;SULF1;HSPB11* |
| regulation of cell differentiation | GO:0045595 | 7/156 | 2.45E-04 | 0.029 | 6.1 | 50.9 | *TFAP2A;TFAP2B;SLC6A6;ZFHX3;CDK6;RUNX3;FGFR2* |
| regulation of filopodium assembly | GO:0051489 | 4/41 | 3.02E-04 | 0.032 | 13.9 | 112.7 | *MYO10;PIK3R1;ARAP1;AGRN* |
| embryonic digestive tract development | GO:0048566 | 3/20 | 5.00E-04 | 0.048 | 22.6 | 171.6 | *PDGFRA;GLI3;FGFR2* |
| neuromuscular junction development | GO:0007528 | 3/24 | 8.67E-04 | 0.058 | 18.3 | 128.8 | *MUSK;LRP4;AGRN* |
| inner ear morphogenesis | GO:0042472 | 3/24 | 8.67E-04 | 0.058 | 18.3 | 128.8 | *TFAP2A;TBX18;FGFR2* |
| positive regulation of phospholipase activity | GO:0010518 | 3/24 | 8.67E-04 | 0.058 | 18.3 | 128.8 | *PDGFRA;ESR1;FGFR2* |
| cellular response to iron ion | GO:0071281 | 2/6 | 9.11E-04 | 0.058 | 63.6 | 445.1 | *TFAP2A;BMP6* |
| positive regulation of hemoglobin biosynthetic process | GO:0046985 | 2/6 | 9.11E-04 | 0.058 | 63.6 | 445.1 | *ABCB10;SLC6A9* |
| positive regulation of filopodium assembly | GO:0051491 | 3/26 | 0.001 | 0.066 | 16.7 | 113.6 | *PIK3R1;ARAP1;AGRN* |
| regulation of hemoglobin biosynthetic process | GO:0046984 | 2/7 | 0.001 | 0.067 | 50.9 | 339.2 | *ABCB10;SLC6A9* |
| sinoatrial node development | GO:0003163 | 2/7 | 0.001 | 0.067 | 50.9 | 339.2 | *TBX18;CACNA1G* |
| receptor clustering | GO:0043113 | 3/28 | 0.001 | 0.069 | 15.3 | 101.1 | *MUSK;LRP4;AGRN* |
| peptidyl-proline hydroxylation to 4-hydroxy-L-proline | GO:0018401 | 2/8 | 0.002 | 0.077 | 42.4 | 270.7 | *P4HA2;P4HTM* |
| negative regulation of alpha-beta T cell differentiation | GO:0046639 | 2/8 | 0.002 | 0.077 | 42.4 | 270.7 | *RUNX3;GLI3* |
| embryonic skeletal system morphogenesis | GO:0048704 | 3/31 | 0.002 | 0.077 | 13.7 | 86.2 | *TFAP2A;PDGFRA;FGFR2* |
| renal water homeostasis | GO:0003091 | 3/31 | 0.002 | 0.077 | 13.7 | 86.2 | *TFAP2B;PRKAR1B;ADCY8* |
| embryonic digestive tract morphogenesis | GO:0048557 | 2/9 | 0.002 | 0.086 | 36.3 | 223.1 | *PDGFRA;FGFR2* |
| negative regulation of epithelial cell proliferation | GO:0050680 | 4/72 | 0.003 | 0.091 | 7.6 | 45.1 | *CDK6;SULF1;RUNX3;FGFR2* |
| cell morphogenesis involved in differentiation | GO:0000904 | 4/72 | 0.003 | 0.091 | 7.6 | 45.1 | *ATRN;LAMA5;ACTN1;PIK3R1* |
| regulation of muscle cell differentiation | GO:0051147 | 3/35 | 0.003 | 0.091 | 12.0 | 71.2 | *CDH2;BOC;FGFR2* |
| forelimb morphogenesis | GO:0035136 | 2/10 | 0.003 | 0.091 | 31.8 | 188.3 | *TFAP2A;TFAP2B* |
| axonogenesis | GO:0007409 | 7/240 | 0.003 | 0.100 | 3.9 | 22.6 | *LAMA5;LAMA2;LAMA3;PIK3R1;S100B;SPTBN1;FGFR2* |
| peptidyl-proline hydroxylation | GO:0019511 | 2/11 | 0.003 | 0.104 | 28.3 | 161.8 | *P4HA2;P4HTM* |
| bone development | GO:0060348 | 3/40 | 0.004 | 0.113 | 10.4 | 57.6 | *TFAP2A;SULF1;FGFR2* |
| regulation of sodium ion transmembrane transporter activity | GO:2000649 | 3/40 | 0.004 | 0.113 | 10.4 | 57.6 | *NETO1;DMD;PTPN3* |
| morphogenesis of a polarized epithelium | GO:0001738 | 2/12 | 0.004 | 0.113 | 25.4 | 141.1 | *LAMA5;LAMA3* |
| skeletal system morphogenesis | GO:0048705 | 3/42 | 0.004 | 0.125 | 9.8 | 53.3 | *TFAP2A;PDGFRA;FGFR2* |
| response to iron ion | GO:0010039 | 2/13 | 0.005 | 0.125 | 23.1 | 124.5 | *TFAP2A;BMP6* |
| bone morphogenesis | GO:0060349 | 2/14 | 0.005 | 0.137 | 21.2 | 111.0 | *TFAP2A;FGFR2* |
| positive regulation of alpha-beta T cell differentiation | GO:0046638 | 2/14 | 0.005 | 0.137 | 21.2 | 111.0 | *RUNX3;GLI3* |
| regulation of phospholipase C activity | GO:1900274 | 2/15 | 0.006 | 0.149 | 19.6 | 99.8 | *PDGFRA;ESR1* |
| response to muscle stretch | GO:0035994 | 2/15 | 0.006 | 0.149 | 19.6 | 99.8 | *CDH2;DMD* |
| regulation of epithelial cell proliferation | GO:0050678 | 4/93 | 0.006 | 0.150 | 5.8 | 29.2 | *CDK6;RUNX3;BMP6;FGFR2* |
| substrate adhesion-dependent cell spreading | GO:0034446 | 3/48 | 0.006 | 0.150 | 8.5 | 42.9 | *LAMA5;ATRN;PIK3R1* |
| hemopoiesis | GO:0030097 | 4/94 | 0.007 | 0.150 | 5.7 | 28.6 | *MEIS1;TOX;RUNX3;FGFR2* |
| digestive tract morphogenesis | GO:0048546 | 2/16 | 0.007 | 0.154 | 18.2 | 90.3 | *PDGFRA;FGFR2* |
| regulation of cellular component movement | GO:0051270 | 3/50 | 0.007 | 0.157 | 8.2 | 40.2 | *CDK6;ACTN1;ARAP1* |
| activation of protein kinase A activity | GO:0034199 | 2/17 | 0.008 | 0.159 | 16.9 | 82.2 | *PRKAR1B;ADCY8* |
| regulation of cellular response to growth factor stimulus | GO:0090287 | 2/17 | 0.008 | 0.159 | 16.9 | 82.2 | *TFAP2B;SULF1* |
| gliogenesis | GO:0042063 | 2/17 | 0.008 | 0.159 | 16.9 | 82.2 | *CDK6;CDH2* |
| synaptic transmission, glutamatergic | GO:0035249 | 2/18 | 0.009 | 0.174 | 15.9 | 75.3 | *GRIK5;GRIK4* |
| cellular response to glucagon stimulus | GO:0071377 | 2/19 | 0.010 | 0.186 | 15.0 | 69.3 | *PRKAR1B;ADCY8* |
| postsynaptic membrane organization | GO:0001941 | 2/19 | 0.010 | 0.186 | 15.0 | 69.3 | *MUSK;LRP4* |
| cellular response to peptide hormone stimulus | GO:0071375 | 4/106 | 0.010 | 0.187 | 5.0 | 23.2 | *PRKAR1B;PIK3R1;ADCY8;CRHR1* |
| chordate embryonic development | GO:0043009 | 3/58 | 0.011 | 0.200 | 7.0 | 31.5 | *PTPRR;SULF1;FGFR2* |
| response to glucagon | GO:0033762 | 2/21 | 0.012 | 0.213 | 13.4 | 59.4 | *PRKAR1B;ADCY8* |
| positive regulation of kinase activity | GO:0033674 | 4/114 | 0.013 | 0.217 | 4.7 | 20.3 | *PDGFRA;MUSK;AGAP2;FGFR2* |
| negative regulation of myeloid cell differentiation | GO:0045638 | 2/22 | 0.013 | 0.217 | 12.7 | 55.3 | *MEIS1;CDK6* |
| regulation of voltage-gated calcium channel activity | GO:1901385 | 2/22 | 0.013 | 0.217 | 12.7 | 55.3 | *DMD;CRHR1* |
| limb morphogenesis | GO:0035108 | 2/22 | 0.013 | 0.217 | 12.7 | 55.3 | *TFAP2B;GLI3* |
| negative regulation of reactive oxygen species metabolic process | GO:2000378 | 2/23 | 0.014 | 0.228 | 12.1 | 51.6 | *TFAP2A;ABCB7* |
| ear morphogenesis | GO:0042471 | 2/23 | 0.014 | 0.228 | 12.1 | 51.6 | *TFAP2A;FGFR2* |
| amino acid import across plasma membrane | GO:0089718 | 2/24 | 0.015 | 0.232 | 11.6 | 48.3 | *SLC6A6;SLC6A9* |
| negative regulation of stress fiber assembly | GO:0051497 | 2/24 | 0.015 | 0.232 | 11.6 | 48.3 | *ARAP1;PIK3R1* |
| positive regulation of neuron apoptotic process | GO:0043525 | 2/24 | 0.015 | 0.232 | 11.6 | 48.3 | *TFAP2A;TFAP2B* |
| lipid translocation | GO:0034204 | 2/24 | 0.015 | 0.232 | 11.6 | 48.3 | *ATP8B1;ATP11A* |
| cardiac conduction system development | GO:0003161 | 2/25 | 0.017 | 0.243 | 11.0 | 45.3 | *TBX18;CACNA1G* |
| in utero embryonic development | GO:0001701 | 2/25 | 0.017 | 0.243 | 11.0 | 45.3 | *PTPRR;FGFR2* |
| positive regulation of ion transmembrane transporter activity | GO:0032414 | 2/26 | 0.018 | 0.250 | 10.6 | 42.6 | *SLC6A9;DMD* |
| synapse organization | GO:0050808 | 4/126 | 0.018 | 0.250 | 4.2 | 16.9 | *CDH2;MUSK;LRP4;AGRN* |
| negative regulation of cation channel activity | GO:2001258 | 2/27 | 0.019 | 0.250 | 10.2 | 40.2 | *SLC6A9;CRHR1* |
| positive regulation of muscle cell differentiation | GO:0051149 | 2/27 | 0.019 | 0.250 | 10.2 | 40.2 | *CDH2;BOC* |
| negative regulation of actin filament bundle assembly | GO:0032232 | 2/27 | 0.019 | 0.250 | 10.2 | 40.2 | *PIK3R1;ARAP1* |
| cell junction organization | GO:0034330 | 3/72 | 0.019 | 0.250 | 5.5 | 21.9 | *LRP4;AGRN;LIMS2* |
| central nervous system development | GO:0007417 | 6/268 | 0.020 | 0.250 | 2.9 | 11.6 | *ZFHX3;MEIS1;CDH2;HESX1;S100B;NPAS1* |
| neutral amino acid transport | GO:0015804 | 2/28 | 0.020 | 0.250 | 9.8 | 38.0 | *SLC6A6;SLC6A9* |
| positive regulation of fibroblast proliferation | GO:0048146 | 2/28 | 0.020 | 0.250 | 9.8 | 38.0 | *PDGFRA;CDK6* |
| positive regulation of phosphatidylinositol 3-kinase activity | GO:0043552 | 2/28 | 0.020 | 0.250 | 9.8 | 38.0 | *PDGFRA;AGAP2* |
| inorganic cation transmembrane transport | GO:0098662 | 6/274 | 0.022 | 0.250 | 2.9 | 11.0 | *SLC6A6;KCNG3;SLC6A8;SLC6A9;ABCB7;CACNA1G* |
| limb development | GO:0060173 | 2/29 | 0.022 | 0.250 | 9.4 | 36.0 | *LRP4;GLI3* |
| regulation of BMP signaling pathway | GO:0030510 | 3/76 | 0.022 | 0.250 | 5.2 | 19.9 | *TFAP2B;SMURF2;SULF1* |
| cell morphogenesis involved in neuron differentiation | GO:0048667 | 3/76 | 0.022 | 0.250 | 5.2 | 19.9 | *LRP4;S100B;FGFR2* |
| axon guidance | GO:0007411 | 5/203 | 0.023 | 0.250 | 3.2 | 12.3 | *LAMA5;LAMA2;LAMA3;PIK3R1;SPTBN1* |
| cardiac muscle cell action potential | GO:0086001 | 2/30 | 0.023 | 0.250 | 9.1 | 34.1 | *DMD;CACNA1G* |
| maintenance of blood-brain barrier | GO:0035633 | 2/30 | 0.023 | 0.250 | 9.1 | 34.1 | *LAMA2;DMD* |
| phospholipid translocation | GO:0045332 | 2/31 | 0.025 | 0.250 | 8.8 | 32.4 | *ATP8B1;ATP11A* |
| regulation of transmembrane receptor protein serine/threonine kinase signaling pathway | GO:0090092 | 2/31 | 0.025 | 0.250 | 8.8 | 32.4 | *TFAP2B;SMURF2* |
| negative regulation of cell cycle | GO:0045786 | 3/80 | 0.026 | 0.250 | 5.0 | 18.2 | *CDK6;RUNX3;PTPN3* |
| chondrocyte differentiation | GO:0002062 | 2/32 | 0.026 | 0.250 | 8.5 | 30.8 | *SULF1;RUNX3* |
| ERBB2 signaling pathway | GO:0038128 | 2/32 | 0.026 | 0.250 | 8.5 | 30.8 | *PTPRR;PIK3R1* |
| regulation of osteoblast differentiation | GO:0045667 | 3/83 | 0.028 | 0.250 | 4.8 | 17.1 | *CDK6;BMP6;FGFR2* |
| positive regulation of plasma membrane bounded cell projection assembly | GO:0120034 | 3/83 | 0.028 | 0.250 | 4.8 | 17.1 | *ARAP1;PIK3R1;AGRN* |
| cell-substrate junction assembly | GO:0007044 | 2/34 | 0.029 | 0.250 | 7.9 | 28.0 | *ACTN1;LAMA3* |
| glucose homeostasis | GO:0042593 | 3/86 | 0.031 | 0.250 | 4.6 | 16.0 | *TFAP2B;PIK3R1;ADCY8* |
| regulation of phosphatidylinositol 3-kinase activity | GO:0043551 | 2/35 | 0.031 | 0.250 | 7.7 | 26.7 | *PDGFRA;PIK3R1* |
| embryonic skeletal system development | GO:0048706 | 2/35 | 0.031 | 0.250 | 7.7 | 26.7 | *PDGFRA;SULF1* |
| regulation of trans-synaptic signaling | GO:0099177 | 2/35 | 0.031 | 0.250 | 7.7 | 26.7 | *GRIK5;GRIK4* |
| muscle cell differentiation | GO:0042692 | 2/35 | 0.031 | 0.250 | 7.7 | 26.7 | *DMD;TBX18* |
| brain development | GO:0007420 | 4/150 | 0.031 | 0.250 | 3.5 | 12.1 | *ZFHX3;MEIS1;CDH2;HESX1* |
| sodium ion transmembrane transport | GO:0035725 | 3/87 | 0.032 | 0.250 | 4.6 | 15.7 | *SLC6A6;SLC6A8;SLC6A9* |
| activation of adenylate cyclase activity | GO:0007190 | 2/36 | 0.033 | 0.250 | 7.5 | 25.5 | *ADCY8;CRHR1* |
| embryonic limb morphogenesis | GO:0030326 | 2/36 | 0.033 | 0.250 | 7.5 | 25.5 | *TFAP2A;GLI3* |
| platelet aggregation | GO:0070527 | 2/36 | 0.033 | 0.250 | 7.5 | 25.5 | *PDGFRA;ACTN1* |
| positive regulation of bone mineralization | GO:0030501 | 2/36 | 0.033 | 0.250 | 7.5 | 25.5 | *TFAP2A;BMP6* |
| membrane assembly | GO:0071709 | 2/36 | 0.033 | 0.250 | 7.5 | 25.5 | *LRP4;SPTBN1* |
| positive regulation of Wnt signaling pathway | GO:0030177 | 4/153 | 0.033 | 0.250 | 3.4 | 11.7 | *SMURF2;AXIN2;SULF1;FGFR2* |
| regulation of erythrocyte differentiation | GO:0045646 | 2/37 | 0.034 | 0.250 | 7.3 | 24.4 | *ABCB10;CDK6* |
| sodium ion transport | GO:0006814 | 3/90 | 0.034 | 0.250 | 4.4 | 14.8 | *SLC6A6;SLC6A8;SLC6A9* |
| positive regulation of cell population proliferation | GO:0008284 | 8/474 | 0.035 | 0.250 | 2.2 | 7.4 | *PDGFRA;TFAP2B;MEIS1;CDK6;TOX;S100B;BMP6;FGFR2* |
| positive regulation of transmembrane receptor protein serine/threonine kinase signaling pathway | GO:0090100 | 3/92 | 0.036 | 0.250 | 4.3 | 14.2 | *SULF1;BMP6;BMP15* |
| odontogenesis | GO:0042476 | 2/39 | 0.038 | 0.250 | 6.9 | 22.4 | *AXIN2;FGFR2* |
| regulation of kinase activity | GO:0043549 | 3/94 | 0.038 | 0.250 | 4.2 | 13.7 | *PDGFRA;MUSK;FGFR2* |
| protein processing involved in protein targeting to mitochondrion | GO:0006627 | 1/5 | 0.039 | 0.250 | 31.6 | 102.6 | *IMMP2L* |
| negative regulation of CD4-positive, alpha-beta T cell differentiation | GO:0043371 | 1/5 | 0.039 | 0.250 | 31.6 | 102.6 | *RUNX3* |
| regulation of adherens junction organization | GO:1903391 | 1/5 | 0.039 | 0.250 | 31.6 | 102.6 | *BMP6* |
| regulation of chloride transport | GO:2001225 | 1/5 | 0.039 | 0.250 | 31.6 | 102.6 | *ATP8B1* |
| regulation of homophilic cell adhesion | GO:1903385 | 1/5 | 0.039 | 0.250 | 31.6 | 102.6 | *PTPRR* |
| cerebral cortex neuron differentiation | GO:0021895 | 1/5 | 0.039 | 0.250 | 31.6 | 102.6 | *TOX* |
| neuroinflammatory response | GO:0150076 | 1/5 | 0.039 | 0.250 | 31.6 | 102.6 | *ADCY8* |
| regulation of microvillus assembly | GO:0032534 | 1/5 | 0.039 | 0.250 | 31.6 | 102.6 | *ATP8B1* |
| regulation of morphogenesis of a branching structure | GO:0060688 | 1/5 | 0.039 | 0.250 | 31.6 | 102.6 | *FGFR2* |
| peripheral nervous system neuron differentiation | GO:0048934 | 1/5 | 0.039 | 0.250 | 31.6 | 102.6 | *RUNX3* |
| positive regulation of CD8-positive, alpha-beta T cell differentiation | GO:0043378 | 1/5 | 0.039 | 0.250 | 31.6 | 102.6 | *RUNX3* |
| G protein-coupled opioid receptor signaling pathway | GO:0038003 | 1/5 | 0.039 | 0.250 | 31.6 | 102.6 | *ADCY8* |
| gastro-intestinal system smooth muscle contraction | GO:0014831 | 1/5 | 0.039 | 0.250 | 31.6 | 102.6 | *SULF1* |
| glial cell-derived neurotrophic factor receptor signaling pathway | GO:0035860 | 1/5 | 0.039 | 0.250 | 31.6 | 102.6 | *SULF1* |
| positive regulation of DNA demethylation | GO:1901537 | 1/5 | 0.039 | 0.250 | 31.6 | 102.6 | *TOX* |
| response to morphine | GO:0043278 | 1/5 | 0.039 | 0.250 | 31.6 | 102.6 | *ADCY8* |
| retina vasculature development in camera-type eye | GO:0061298 | 1/5 | 0.039 | 0.250 | 31.6 | 102.6 | *PDGFRA* |
| positive regulation of long-term synaptic depression | GO:1900454 | 1/5 | 0.039 | 0.250 | 31.6 | 102.6 | *ADCY8* |
| SA node cell to atrial cardiac muscle cell signaling | GO:0086018 | 1/5 | 0.039 | 0.250 | 31.6 | 102.6 | *CACNA1G* |
| salivary gland morphogenesis | GO:0007435 | 1/5 | 0.039 | 0.250 | 31.6 | 102.6 | *FGFR2* |
| positive regulation of necrotic cell death | GO:0010940 | 1/5 | 0.039 | 0.250 | 31.6 | 102.6 | *SLC6A6* |
| sinoatrial node cell differentiation | GO:0060921 | 1/5 | 0.039 | 0.250 | 31.6 | 102.6 | *TBX18* |
| membrane depolarization during AV node cell action potential | GO:0086045 | 1/5 | 0.039 | 0.250 | 31.6 | 102.6 | *CACNA1G* |
| membrane depolarization during SA node cell action potential | GO:0086046 | 1/5 | 0.039 | 0.250 | 31.6 | 102.6 | *CACNA1G* |
| trivalent inorganic anion homeostasis | GO:0072506 | 1/5 | 0.039 | 0.250 | 31.6 | 102.6 | *TFAP2B* |
| ubiquitin-dependent SMAD protein catabolic process | GO:0030579 | 1/5 | 0.039 | 0.250 | 31.6 | 102.6 | *SMURF2* |
| positive regulation of protein kinase B signaling | GO:0051897 | 4/161 | 0.039 | 0.250 | 3.3 | 10.6 | *PDGFRA;PIK3R1;ESR1;FGFR2* |
| digestive tract development | GO:0048565 | 2/40 | 0.040 | 0.250 | 6.7 | 21.5 | *GLI3;FGFR2* |
| positive regulation of ossification | GO:0045778 | 2/40 | 0.040 | 0.250 | 6.7 | 21.5 | *TFAP2A;BMP6* |
| positive regulation of biomineral tissue development | GO:0070169 | 2/41 | 0.042 | 0.250 | 6.5 | 20.7 | *TFAP2A;BMP6* |
| negative regulation of canonical Wnt signaling pathway | GO:0090090 | 4/165 | 0.042 | 0.250 | 3.2 | 10.1 | *LRP4;AXIN2;GLI3;TBX18* |
| regulation of neuron apoptotic process | GO:0043523 | 3/98 | 0.043 | 0.250 | 4.0 | 12.7 | *TFAP2A;TFAP2B;AGAP2* |
| negative regulation of protein catabolic process | GO:0042177 | 2/42 | 0.043 | 0.250 | 6.3 | 19.9 | *AGAP2;PTPN3* |
| negative regulation of transcription by RNA polymerase II | GO:0000122 | 10/684 | 0.045 | 0.250 | 1.9 | 5.9 | *TFAP2A;TFAP2B;ZFHX3;SMURF2;HESX1;RUNX3;ESR1;BMP6;FGFR2;GLI3* |
| cellular glucose homeostasis | GO:0001678 | 2/43 | 0.045 | 0.250 | 6.2 | 19.2 | *PIK3R1;ADCY8* |
| positive regulation of phospholipase C activity | GO:0010863 | 2/43 | 0.045 | 0.250 | 6.2 | 19.2 | *PDGFRA;ESR1* |
| adherens junction maintenance | GO:0034334 | 1/6 | 0.046 | 0.250 | 25.3 | 77.5 | *KIFC3* |
| astrocyte development | GO:0014002 | 1/6 | 0.046 | 0.250 | 25.3 | 77.5 | *CDK6* |
| negative regulation of morphogenesis of an epithelium | GO:1905331 | 1/6 | 0.046 | 0.250 | 25.3 | 77.5 | *SULF1* |
| regulation of DNA demethylation | GO:1901535 | 1/6 | 0.046 | 0.250 | 25.3 | 77.5 | *TOX* |
| regulation of focal adhesion disassembly | GO:0120182 | 1/6 | 0.046 | 0.250 | 25.3 | 77.5 | *PIK3R1* |
| nephron tubule development | GO:0072080 | 1/6 | 0.046 | 0.250 | 25.3 | 77.5 | *TFAP2B* |
| chondrocyte development | GO:0002063 | 1/6 | 0.046 | 0.250 | 25.3 | 77.5 | *SULF1* |
| peripheral nervous system neuron development | GO:0048935 | 1/6 | 0.046 | 0.250 | 25.3 | 77.5 | *RUNX3* |
| regulation of peptidyl-cysteine S-nitrosylation | GO:2000169 | 1/6 | 0.046 | 0.250 | 25.3 | 77.5 | *DMD* |
| embryonic camera-type eye morphogenesis | GO:0048596 | 1/6 | 0.046 | 0.250 | 25.3 | 77.5 | *TFAP2A* |
| epithelial tube branching involved in lung morphogenesis | GO:0060441 | 1/6 | 0.046 | 0.250 | 25.3 | 77.5 | *FGFR2* |
| epithelial tube formation | GO:0072175 | 1/6 | 0.046 | 0.250 | 25.3 | 77.5 | *FGFR2* |
| regulation of tooth mineralization | GO:0070170 | 1/6 | 0.046 | 0.250 | 25.3 | 77.5 | *TFAP2A* |
| gamma-aminobutyric acid transport | GO:0015812 | 1/6 | 0.046 | 0.250 | 25.3 | 77.5 | *SLC6A6* |
| glandular epithelial cell differentiation | GO:0002067 | 1/6 | 0.046 | 0.250 | 25.3 | 77.5 | *FGFR2* |
| positive regulation of focal adhesion disassembly | GO:0120183 | 1/6 | 0.046 | 0.250 | 25.3 | 77.5 | *PIK3R1* |
| inner ear auditory receptor cell differentiation | GO:0042491 | 1/6 | 0.046 | 0.250 | 25.3 | 77.5 | *SLC4A7* |
| sialylation | GO:0097503 | 1/6 | 0.046 | 0.250 | 25.3 | 77.5 | *ST6GAL1* |
| magnesium ion homeostasis | GO:0010960 | 1/6 | 0.046 | 0.250 | 25.3 | 77.5 | *TFAP2B* |
| maintenance of DNA repeat elements | GO:0043570 | 1/6 | 0.046 | 0.250 | 25.3 | 77.5 | *AXIN2* |
| trigeminal nerve development | GO:0021559 | 1/6 | 0.046 | 0.250 | 25.3 | 77.5 | *TFAP2A* |
| positive regulation of tooth mineralization | GO:0070172 | 1/6 | 0.046 | 0.250 | 25.3 | 77.5 | *TFAP2A* |
| neurogenesis | GO:0022008 | 2/44 | 0.047 | 0.250 | 6.0 | 18.4 | *CDK6;BTBD6* |
| embryonic organ morphogenesis | GO:0048562 | 2/44 | 0.047 | 0.250 | 6.0 | 18.4 | *PDGFRA;FGFR2* |
| homotypic cell-cell adhesion | GO:0034109 | 2/44 | 0.047 | 0.250 | 6.0 | 18.4 | *PDGFRA;ACTN1* |
| action potential | GO:0001508 | 2/45 | 0.049 | 0.250 | 5.9 | 17.8 | *DMD;CACNA1G* |
| positive regulation of signal transduction | GO:0009967 | 5/252 | 0.050 | 0.250 | 2.6 | 7.8 | *PIK3R1;SULF1;BMP6;FGFR2;LIMS2* |

Note. The full enrichment analysis for genes shared between fish early domesticates and human groups with autism spectrum disorders can be accessed here: <https://maayanlab.cloud/Enrichr/enrich?dataset=c5b8b850d070c8fa90b809675170affd>
